# Pleiotropic Influence of DNA Methylation QTLs on physiological and aging traits

**DOI:** 10.1101/2023.04.12.536608

**Authors:** Khyobeni Mozhui, Hyeonju Kim, Flavia Villani, Amin Haghani, Saunak Sen, Steve Horvath

## Abstract

DNA methylation is influenced by genetic and non-genetic factors. Here, we chart quantitative trait loci (QTLs) that modulate levels of methylation at highly conserved CpGs using liver methylome data from mouse strains belonging to the BXD Family. A regulatory hotspot on chromosome 5 had the highest density of trans-acting methylation QTLs (trans-meQTLs) associated with multiple distant CpGs. We refer to this locus as meQTL.5a. The trans-modulated CpGs showed age-dependent changes, and were enriched in developmental genes, including several members of the MODY pathway (maturity onset diabetes of the young). The joint modulation by genotype and aging resulted in a more “aged methylome” for BXD strains that inherited the DBA/2J parental allele at meQTL.5a. Further, several gene expression traits, body weight, and lipid levels mapped to meQTL.5a, and there was a modest linkage with lifespan. DNA binding motif and protein-protein interaction enrichment analysis identified the hepatic nuclear factor, *Hnf1a* (MODY3 gene in humans), as a strong candidate. The pleiotropic effects of meQTL.5a could contribute to variation in body size and metabolic traits, and influence CpG methylation and epigenetic aging that could have an impact on lifespan.

## Introduction

Genome-wide patterns in DNA methylation (DNAm) are established during development and are critical for cell differentiation and cell identity.^1^ The canonical form of DNAm involves the addition of a methyl-group to the cytosine residue at CG dinucleotides (i.e., CpG methylation). The methylation status of CpGs is a part of the epigenetic landscape that serves as a stable and yet reprogrammable form of gene expression regulation.^2, 3^ On one hand, methylation of CpGs are important for sustaining and perpetuating expression signatures and in giving each organ and tissue its functional identity. On the other hand, the methylome is dynamic and a modifiable molecular process that enables the genome to respond and adapt to ever changing environmental and nutritional states.^4, 5^ Due to its modifiability, DNAm is profoundly altered by the passage of time, and tracks closely with age and aging.^6, 7^

Along with aging and modifications by extrinsic factors, CpG methylation can also be influenced by underlying genetic sequence variants. Several studies in humans have identified genetic loci that are associated with DNAm.^8–10^ Analogues to gene expression quantitative trait loci (eQTLs),^11^ a genetic region that influences the quantitative variation in CpG methylation is referred to as a methylation QTL, or meQTL (alternatively also shortened to “mQTL”; although that can be confused with “metabolite QTL” and “module QTL”^12, 13^). Several human studies have performed genome-wide association studies (GWAS) for CpG methylation, and there are now large-scale multi-tissue meQTL atlases available for humans.^14, 15^ Similar to the classification of eQTLs into cis- and trans-effects, meQTLs are also categorized into cis-meQTLs or trans-meQTLs, depending on the distance between the meQTL regulatory locus, and the target CpG.^15, 16^ Cis-meQTLs are highly enriched for genetic loci that have been associated with complex traits, and genetic variation in methylation levels are implicated in disease risk.^9, 10, 17, 18^ An meQTL region can be associated with multiple distal CpGs in trans, and similar to trans-eQTL hotpots, such sites represent trans-meQTL hotspots.^8, 10, 19^ Trans-meQTL hotspots implicate causal modulators with widespread influence on CpGs. There is now growing evidence that DNA binding factors and transcription factors (TFs) play a role in shaping the methylome by exerting trans-regulatory influence on CpGs.^10, 20–22^ For instance, during hepatocyte differentiation, TFs such as the hepatocyte nuclear factors (HNFs) and GATA family are reported to regulate the dynamic spatial and temporal patterns in DNAm.^23^

Model organisms provide a powerful tool for interrogating the interactions between meQTLs, eQTLs, and experimental conditions such as diets and drugs. However, although genome-scale meQTL studies in humans date back to the 2010s,^24^ there is an over 10-year lag in methylome-wide meQTL studies in rodent populations. This is partly due to the lack of a cost-effective and scalable DNAm microarray for model organisms that is comparable to the Illumina HumanMethylation Infinium BeadChips.^25^ A few of us attempted to repurpose the human arrays to measure methylation in mice.^26–28^ Indeed a small proportion of the CpG probes on the human arrays map to conserved sequences, and could be considered as “pan-mammalian” interrogators of the epigenome. More recently, a truly pan-mammal array, the HorvathMammalMethylChip40, was custom developed.^29, 30^ A unique aspect of the array is that the probes map to conserved sequences, and this has opened up new avenues for large multi-species comparative epigenomics.^31, 32^ We used this array to track epigenetic changes with aging when mice are subjected to two different dietary conditions.^6^ Our data incorporated genetic diversity as we profiled members on the BXD Family. In the present work, we use the methylome data for an meQTL mapping study.

The BXDs are a family of recombinant inbred (RI) and advanced intercross (AI) mouse strains. We have previously described the BXDs in greater detail.^33–36^ In brief, the BXD Family consists of about 150 inbred members derived from two progenitor strains: C57BL/6J (B6) and DBA/2J (D2). The BXDs have a long history in quantitative genetics and the earlier sets of RI strains were used to map simpler Mendelian traits.^37, 38^ Subsequently, additional sets of RI and AI strains were added to the growing family, and over the years, the BXDs have accrued a vast compendium of phenotypic data ranging from metabolic, physiologic, lifespan, to behavioral and neural traits, and multi-omic datasets (e.g., transcriptomics, proteomics, and metabolomics).^35, 39–41^ This is matched by deep genome sequence data with over 6 million genetic variants segregating in the family, making the BXDs a powerful mammalian panel for systems genetics, and systems epigenetics.^6, 30, 33, 42, 43^

In our previous work, we studied the genetic regulation of epigenetic clocks in the BXDs and examined metabolic and dietary factors that are related to the age-dependent methylation changes.^6^ Here, we focus on meQTLs that influence individual CpGs, and evaluate the genetic architecture of CpG methylation in liver tissue. We identify meQTL hotspots that influence multiple distal CpGs. The region on chromosome (Chr) 5 harbored the highest density of co-localized trans-meQTLs, and we refer to this genetic interval as meQTL.5a. This region also contains a high-density of QTLs linked to gene expression both at the transcriptomic (eQTLs) and proteomic (pQTLs) levels. For the CpGs that are trans-modulated by meQTL.5a, the pattern of variance indicates a genotype dependent susceptibility to the effects of aging, and to an extent, diet. Specifically, we find a more aged methylome for strains that have the D2 allele at meQTL.5a. Further, we find a pleiotropic effect of this locus on body weight, and for this, the B6 allele was associated with a positive additive effect. The contrasting allelic effects on the two traits may moderate the impact on this locus on lifespan. Based on DNA binding motif enrichment and protein-protein interaction (PPI), we identify the hepatic nuclear factor, *Hnf1a*, as one of the important candidate genes in meQTL.5a. In humans, mutations in *HNF1A* results in MODY3 (maturity onset diabetes of the young 3),^44^ and our results indicate a trans-modulatory effect of meQTL.5a on CpGs located in several other genes that are part of the MODY pathway. Overall, our results suggest a convergent effect of age and diet on CpGs that are also partly influenced by an meQTL. We propose a model in which the meQTL.5a has both a horizontal and vertical pleiotropic effect on physiological traits, DNAm, and lifespan.

## Result

### Overview of meQTL distribution in the mouse genome

The HorvathMammalMethylChip40 array contains probes for 27966 CpGs that have been validated for the mouse genome.^6, 30^ We have used this to assay the liver methylome in a population of BXDs (details on the samples are in **Data S1**). To uncover genetic loci that modulate methylation variation at these CpGs, we performed linkage mapping across the autosomal chromosome (Chrs).^33^ QTL mapping was implemented in R/qtl2 and adjusted for age, diet, the top methylome-wide principal component, and the BXD’s kinship matrix.^45^ For each of the 27966 CpGs, we plotted its genome-wide highest LOD score, and the distance between the maximal meQTL marker, and the CpG location (**Fig 1a**; genome-wide peak LOD data for each CpG is in **Data S2**). Strong meQTLs tended to reside within 10 Mb of the corresponding CpGs, and we classified these regulatory loci as cis-meQTLs, and the targeted CpG as cis-CpGs. Note that due to the family-based population and the comparatively larger haplotypes in the BXDs,^33^ we have used a much larger interval rather than the <1 Mb interval that is typically used to assign cis-effects in human association studies.^8, 15^ In total, 3921 CpGs (14% of the CpGs we examined) mapped to at least one meQTL at a nominal LOD ≥ 3.5. Many of the CpGs were polygenic and mapped to more than one locus (in other words, a CpG with a strong cis-meQTL may also have lower QTLs in trans). The meQTLs showed an uneven distribution with some loci having a trans-modulatory linkage to many distal CpGs that potentially signify a regulatory hotspot (**Fig 1b**). At such trans-meQTL hotspots, there is an imbalance in which parental allele increased methylation (i.e., has the positive additive effect). For instance, majority of the CpGs that have meQTLs on markers on Chr 5 (∼115 Mb) are associated with higher methylation for the allele from the D2 parent (*D* allele) (**Fig 1b**). On Chr2 (∼110 Mb), it is the allele from the B6 parent (*B* allele) that is associated with higher methylation. This is consistent with reports from human studies that SNPs associated with multiple CpGs in trans have the same direction of allelic effect.^22^ The allelic effects are clearly visible when we consider only the genome-wide peak LOD markers for each CpG (i.e., each CpG linked to only its genome-wide strongest meQTL marker). **Fig 1c** plots the locations of 1416 peak LOD markers against the location of 3921 CpGs that mapped at LOD ≥ 3.5. Of these, 1833 CpGs mapped as cis-meQTLs (meQTLs to marker ratio of 1.98). The remaining 2088 CpGs had peak LOD at 691 unique markers that were distant from the location of the CpG (meQTL to marker ratio of 3.02). These QTLs are classified as trans-meQTLs, and the CpGs are referred to as trans-CpGs. For the cis-meQTLs, the number of loci in which the *B* allele had the positive addictive effect (909 cis-meQTLs) was similar to the number of loci in which the *D* allele had the positive additive effect (924 cis-meQTLs). However, for the trans-meQTLs, there was a preponderance for higher methylation for the *D* allele (1413 or 68% of the trans-CpGs).

**Fig 1.**
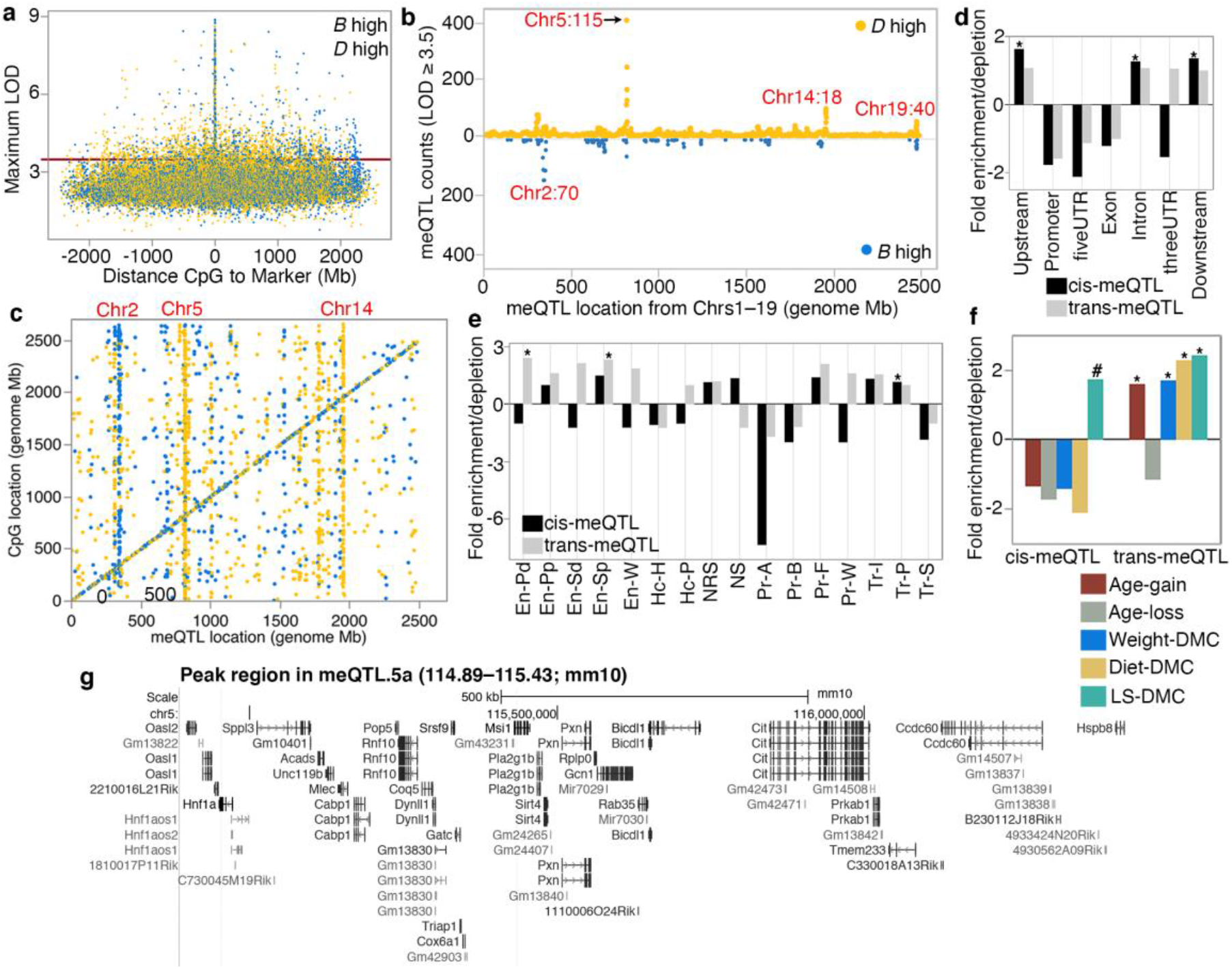
Overview of methylation QTL (meQTLs) in the liver. **(a)** Plot of the genome-wide peak LOD score for the 27966 CpGs, and distance between the CpG and the maximum LOD marker. Yellow: *D* allele (DBA/2J genotype) has positive additive effect; blue: *B* allele (C57BL/6J genotype) has positive additive effect. **(b)** meQTL location (x-axis; from chromosomes 1–19) and counts of meQTLs with LOD ≥ 3.5. **(c)** The genome graph plots location of the genome-wide peak QTL marker (x-axis), and location of the linked CpG (y-axis). Shows only the 3921 CpGs that map at LOD ≥ 3.5. Relative enrichment or depletion in **(d)** predicted chromatin states, and **(e)** genomic location for CpGs that map as cis-meQTLs (black), and as trans-meQTLs (grey). Asterisks denote hypergeometric enrichment p < 0.001. **(f)** Enrichment or depletion in differentially methylated CpGs among the cis- and trans-modulated CpGs. (Asterisks denote hypergeometric enrichment p < 0.001; hash denotes p = 0.003) **(g)** Portion of the peak interval in the chromosome 5 meQTL hotspot: meQTL.5a (from UCSC Genome browser GRCm38/mm10).

In terms of genomic locations and chromatin states, the cis-CpGs were enriched for introns and intergenic regions, and were located in transcriptionally permissive states (Tr-P), but were highly depleted in active and bivalent promoters (Pr-A and Pr-B, respectively), transcriptionally strong states (Tr-S) (**Fig 1d,e**; **Table S1**), and gene exons, promoters and 3’ and 5’ UTRs. Trans-CpGs on the other hand, were enriched in enhancer sites, and depleted in promoter regions (**Fig 1d, e**; **Table S1**).

To examine how the genetic variation in methylation relate to variance associated with aging, diet, body weight, and genotype dependent longevity (variables that we have reported in detail in ^6^), we examined the proportion of differentially methylated CpGs (DMCs) that map as cis- or trans-meQTLs. The cis-CpGs were only modestly enriched in DMCs associated with genotype-dependent lifespan (lifespan differentially methylated CpGs or LS-DMC; hypergeometric enrichment p = 0.003), and were depleted in DMCs related to age, weight, and diet (**Fig 1f**; **Table S2**). This indicates that variance of CpGs that are under cis-modulation are largely due to genetics. In contrast, the trans-CpGs were highly enriched in CpGs that gained methylation with age (age-gain), and CpGs associated with weight, diet, and LS-DMCs. This suggests that variance of CpGs that are under trans-modulation are multi-factorial and influenced by both genetic and non-genetic factors.

### meQTL hotspots and association with gene expression

To define regions that contain a high density of meQTLs, we took the 3921 CpGs with maximal LOD ≥ 3.5 and counted the number of meQTLs linked to each genotype marker. 18 markers were associated with 20 or more meQTLs and we classified these as putative meQTL hotspots (**Table S3**). A few of these are mostly cis-meQTL regions. For example, the two neighboring markers on Chr19, rs30567369 (47.51 Mb) and rs31157694 (47.94 Mb), were linked to 58 CpGs in cis. We have previously reported this region as a QTL for liver epigenetic age acceleration (distal portion of “epigenetic age acceleration QTL on Chr19” or Eaa19).^6^

For trans effects, the highest number of trans-meQTLs per marker was on Chr5, ∼115 Mb (**Fig 1b, c, g**). Here, the SNP marker rs29733222 (115.43 Mb; coordinates based on GRCm38/mm10) is linked to 230 genome-wide peak trans-meQTLs, and only 3 genome-wide peak cis-meQTLs (**Table S3**). As several neighboring markers in this Chr5 region were linked to multiple meQTLs, we roughly delineated a 10 Mb interval (110–120 Mb) as a liver meQTL hotspot and refer this region as meQTL.5a. In total, 535 meQTLs mapped to meQTL.5a at the 3.5 LOD score threshold (502 trans-meQTLs, 33 cis-meQTLs; **Table 1**; **Data S2**). Majority of the trans-meQTLs in meQTL.5a (435 of the 502) were associated with higher methylation for the *D* allele, and only 67 trans-meQTLs were associated with higher DNAm for the *B* allele.

**Table 1.** Liver meQTL hotspots

| Name | Broad location (peak) | Liver meQTL | Hotspot type | Module QTL | Liver eQTL | me/eQTL overlap |
| --- | --- | --- | --- | --- | --- | --- |
| meQTL.2a | Chr2:102–112 (107 Mb) | 72 (41 cis) | cis methyl |  | 80 (13 cis) | *Them7* (cg10774906); *Mpped2* (cg07667286, cg00811894) |
| meQTL.2b | Chr2:141–151 (144–149 Mb) | 164 (15 cis) | trans methyl | Green (2092) | 27 (13 cis) | *Rem1* (cg23754359, cg25361894); *Rin2* (cg12687767) |
| meQTL.4a | Chr4:114–124 (119 Mb) | 39 cis | cis methyl |  | 43 (25 cis) | *Faah* (cg08815464, cg15610892, cg19641802, cg06184921) |
| meQTL.5a | Chr5:110–120 (115 Mb) | 535 (33 cis) | trans methyl | Blue (5067); Black (1087) | 376 (36 cis) | *Tenm3* (trans; cg24399106), *Clcn3* (trans; cg16842643) |
| meQTL.7a | Chr7:132–142 (138 Mb) | 59 (9 cis) | trans methyl |  | 197 (7 cis) | *Mgmt* (cg00046614, cg11711038, cg23272565) |
| meQTL.14a | Chr14:17–27 (23 Mb) | 51 (32 cis) | cis methyl |  | 5 cis | - |
| meQTL.14b | Chr14: 41–51 (46–49 Mb) | 184 (19 cis) | trans methyl | Greenyellow (998) | 97 (31 cis) | *Ddhd1* (cg17695612; cg10924987); *Prmt5* (cg24106188) |
| distal Eaa19<sup>1</sup> | Chr19:42–52 (48 Mb) | 103 (100 cis) | cis methyl | Royalblue (62); Lightgreen (1761) | 274 (43 cis) | *Crtac1*; *Ldb1*; *Psd*; *Col17a1* |

<sup>1</sup>Epigenetic age acceleration

Similarly, we demarcated broad 10 Mb intervals around the other putative meQTL hotspots and counted the number of CpGs that have peak LOD scores in these intervals. We tabulated 8 putative meQTL hotspots that are in Chrs 2, 4, 5, 7, 14, and 19 (**Table 1**). To examine if these meQTL intervals also influence gene expression we referred to an existing BXD liver RNA-seq data (previously reported in ^41^) and searched for transcripts that map to the 10 Mb intervals listed in **Table 1** at eQTL LOD ≥ 3.5. meQTL.5a had the largest number of eQTLs, followed by Eaa19 (**Table 1**; lists of transcripts that map to meQTL.5a and Eaa19 are in **Data S3** and **Data S4**). meQTL.5a had 376 liver eQTLs that included 36 cis-eQTLs from positional candidate genes such as *Cit*, *Sirt4*, and *Hspb8*. Eaa19, despite being primarily a cis-meQTL locus, had an abundance of trans-eQTLs (**Table 1**). Somewhat surprisingly, for all the meQTL intervals, there was very little overlap between meQTLs and eQTLs, even for the strong cis-effects that suggests limited co-regulation of the methylome and the transcriptome. Only a few genes (listed in **Table 1**) had concordant meQTLs and eQTLs in the same locus, and of these, only the QTLs for *Clcn3* and *Tenm3* in meQTL.5a were *trans-*effects. For both *Clcn3* and *Tenm3*, the trans-modulated CpGs (cg16842643 and cg24399106, respectively) are in the 5’UTR. The remaining few genes with overlapping me/eQTLs were cis-effects.

We prioritized the meQTL.5a and Eaa19 intervals and referred to the liver proteomic data (also reported in ^41^) to search for protein QTLs (pQTLs) in meQTL.5a and Eaa19. At the same LOD ≥ 3.5 threshold, 104 protein variants from 83 unique genes mapped as pQTLs to meQTL.5a (32 cis-pQTLs). There was more consistency between pQTLs and eQTLs, and *Hsd17b4*, *Psmb8*, *Psmb9*, and *Psmb10* had trans-eQTLs and trans-pQTLs, and *Pebp1* had cis-eQTL and cis-pQTL in meQTL.5a (**Data S3**). Similarly, the overlap between eQTLs and pQTLs was higher for Eaa19. In total, 138 protein variants (57 cis) mapped to Eaa19, and of these, *Abcc2*, *Cutc*, *Cyp2c70*, *Gsto*, and *Sfxn2* had cis-acting QTLs for both mRNA and protein, and *Cyp1a1* and *Naga* had trans-eQTLs and trans-pQTLs (**Data S4**).

### Genetic modulation of co-methylation networks in mouse liver

We applied a weighted gene co-methylation network analysis (WGCNA) to evaluate whether the meQTL hotspots could be detected at the network level.^46–49^ WGCNA was carried out on the set of ∼28K CpGs. At a soft-threshold power of 6, the CpGs were grouped into 14 modules that range in size from 62–13821 CpG members that form tightly correlated networks (**Data S5; Fig S1a**; the module membership for each of the CpGs are in **Data S2**). For each module, the module eigengene (ME) is the top principal component of the co-methylation network and is the representative methylation pattern.^46^ The inter-module correlations between the MEs provide a view of the covariance among the CpG networks (i.e., meta-network) (**Data S5** and displayed in **Fig S1b**). The MEs can also be tested for association with major variables such as age and diet, and this is a convenient way to assess the network-level impact of these variables.^49^ Unsurprisingly, age was a significant correlate of the CpG networks, and 6 of the 14 modules were significantly correlated with age (p < 0.001, |r| ≥0.18; **Data S5**). Of these, the Green module (2092 CpG members), followed by the Lightgreen module (1761 CpGs), had the tightest correlation with age (r = 0.69 and 0.49, respectively; **Data S5** and **Fig S1c, S1d**). The large Blue module with 5067 CpG members was significantly anti-correlated with age (r = –0.34).

Our primary focus is on genetic modulation of these CpG networks, and we performed QTL mapping for each of the MEs with age, diet, and body weight as co-factors. The module-level QTL mapping was done using the Genome-wide Efficient Mixed Model (GEMMA) algorithm implemented on the webtool GeneNetwork.^50–52^ The strongest QTL was for the small Royalblue module, which mapped at LOD = 27 to distal Eaa19 (QTL plots for select modules in **Fig 2**; full QTL results in **Data S6** and **Fig S2**). The age-associated modules, Blue and Lightgreen, also had modest QTLs in Eaa19 (**Fig 2**). The large Blue module that is anticorrelated with age mapped at a LOD = 4.6 to meQTL.5a (**Fig 2**). The Black and age-associated Lightgreen modules also had modest QTLs in meQTL.5a (**Data S6; Fig 2**). For the Royalblue module, 45 of the 62 CpGs members were located in Eaa19, and this module mostly represented a correlated network of CpGs that are cis-modulated by variants in Eaa19. In contrast, only 29 of the 5067 CpGs in the Blue module were located in meQTL.5a, and indicates that the Blue ME captures CpGs that have shared covariance due to a trans-effect from meQTL.5a. The Blue module members include 443 of the 502 CpGs that had genome-wide peak trans-meQTLs at LOD ≥ 3.5 in meQTL.5a.

**Fig 2.**
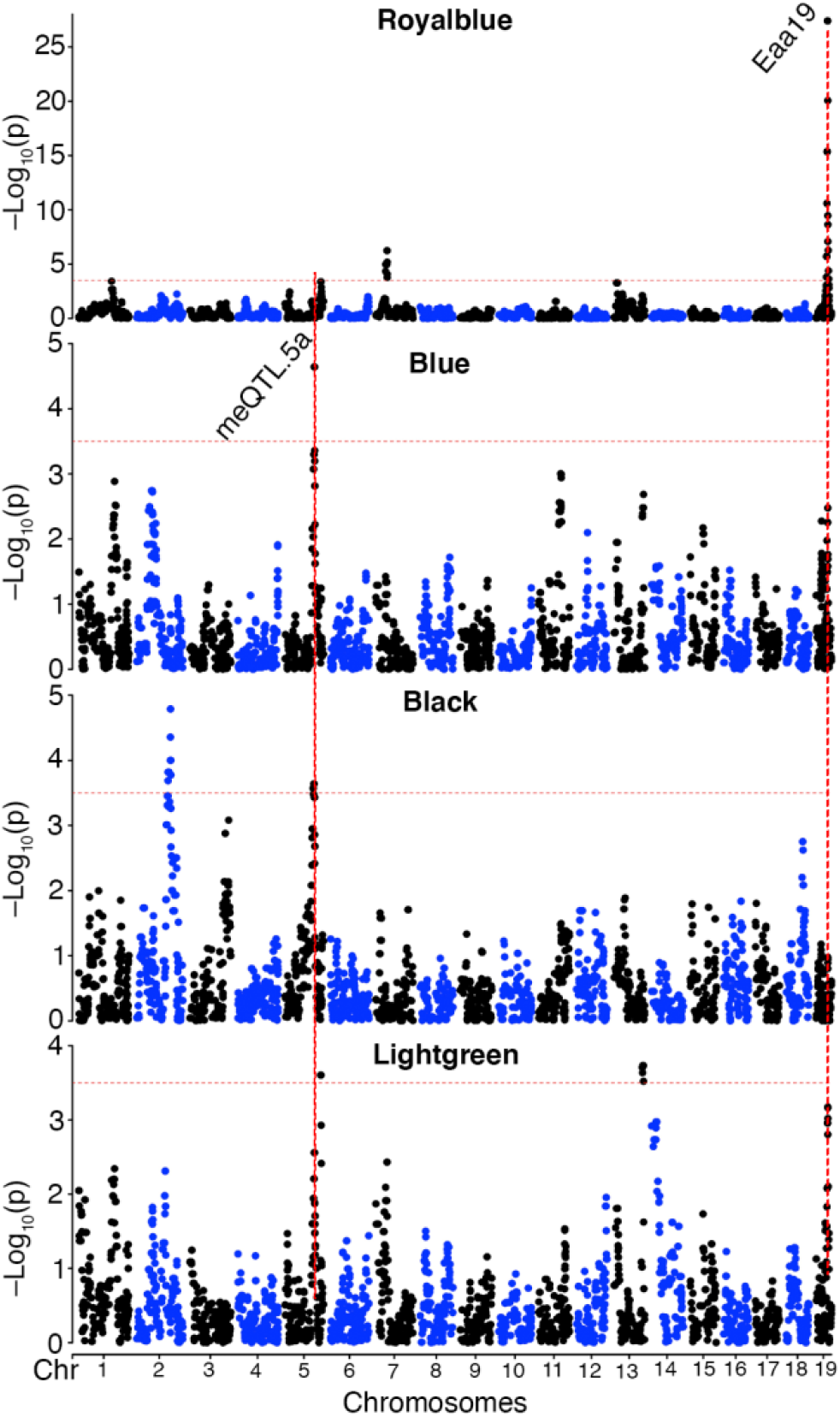
Genetics of co-methylation CpG networks. QTL maps for four module eigengenes (MEs) are shown. Mapping was done using a linear mix model. The horizontal dashed red line marks a relatively lenient threshold of –log_10_p = 3.5. The Blue, Black and Lightgreen modules share suggestive overlapping QTLs in meQTL.5a. Eaa19 has a strong cis-regulatory effect on the Royalblue module. The chromosome 2 peak for the Black module is proximal to meQTL.2b in Table 1.

Overall, the WGCNA shows that multiple distal CpGs can form tightly correlated networks partly due to shared genetic modulation, and once again highlights meQTL.5a as a CpG regulatory hotspot.

### Characterizing the CpGs trans-modulated by meQTL.5a

To uncover common biological pathways among the set of trans-CpGs linked to meQTL.5a, we performed a genomic regions enrichment analysis using the GREAT tool.^53, 54^ Compared to the background array, the 502 trans-CpGs were highly enriched in developmental and cell differentiation genes (**Data S7**; **Fig 3a**). The CpG regions were also enriched in promoter motifs including sequences that are downstream targets of the hepatocyte nuclear factor 4, alpha (HNF4A). Additionally, the FOXA1 (HNF3A) TF network was an enriched pathway among the meQTL.5a trans-CpGs. A regional enrichment analysis for the CpGs in the Blue module highlighted the same pathways and TF networks (**Data S7; Fig 3b**), and this collectively suggests that the CpGs modulated by meQTL.5a are related to development, and may be targeted by related DNA binding factors, particularly the hepatocyte nuclear factors. In terms of genomic context, compared to the background array, the trans-CpGs targeted by meQTL.5a were highly enriched in predicted enhancer states (e.g., En-Pd, En-Pp, En-Sd, En-Sp, and En-W),^6, 55, 56^ and were mostly located in introns (**Table S4**).

**Fig 3.**
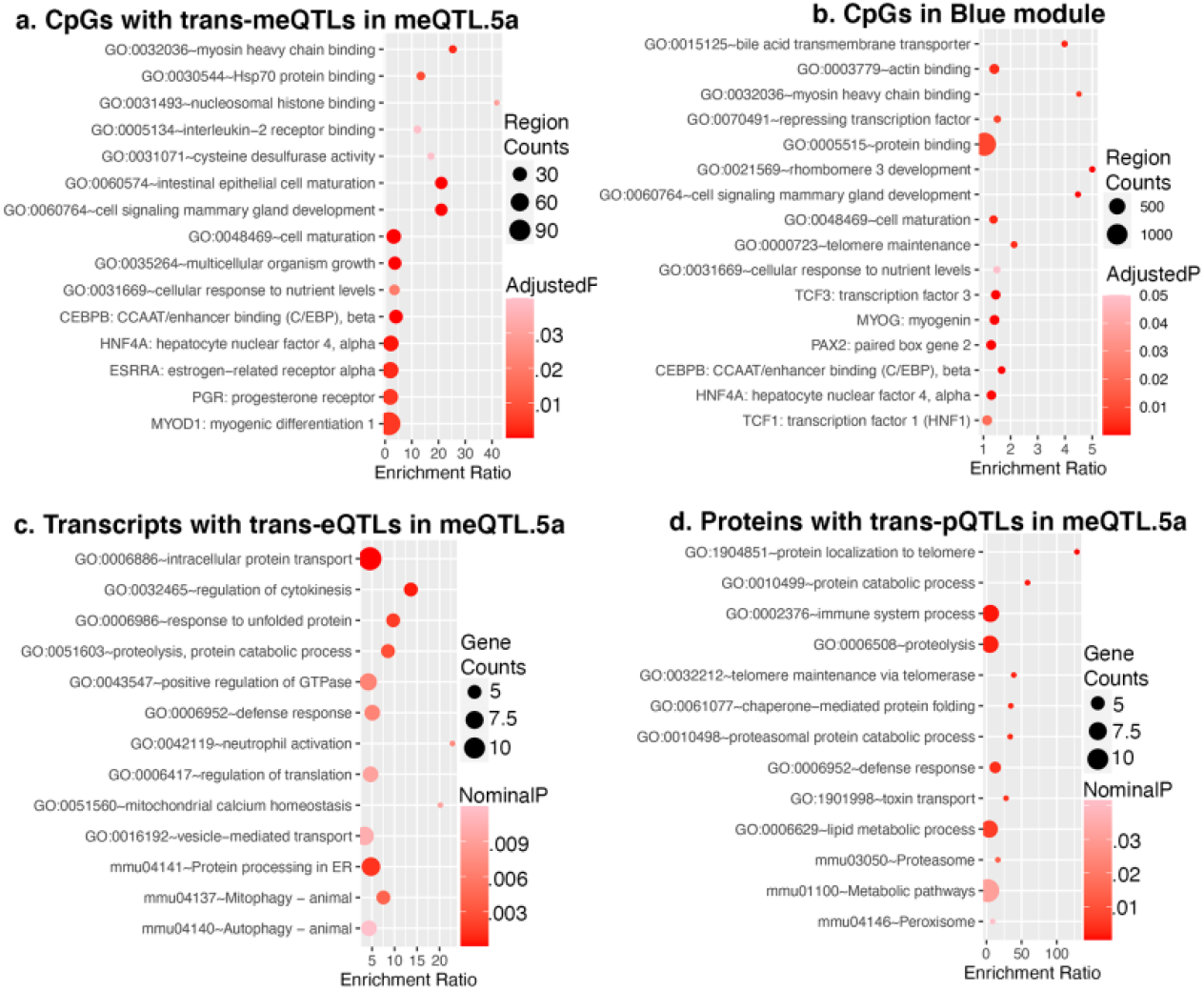
Enriched functional pathways. Genomic regions enrichment analysis of CpGs that are trans-modulated by meQTL.5a **(a)**, and CpGs that are members of the Blue module **(b)**. Gene ontology and pathway enrichment among **(c)** transcripts, and **(d)** proteins that map to meQTL.5a in liver.

To test if we find intersecting biological functions in the transcriptome and proteome, we performed a gene ontology (GO) enrichment analysis of the trans-modulated mRNAs and proteins that map to meQTL.5a (**Data S8**). There were no functionally enriched categories after FDR correction. At a nominal p-value, the top 10 GO (for biological processes), and top KEGG pathways for both the trans-modulated proteins and transcripts were related to protein transport and protein catabolism (**Fig 3b, 3c**; **Data S8**). The trans-pQTLs were also nominally enriched in telomere maintenance, and the trans-eQTLs in mitophagy and autophagy. However, there was limited overlap in functional pathways between the trans-modulated CpGs and trans-modulated mRNA/proteins.

### High-priority candidate genes in meQTL.5a

The peak markers in linkage disequilibrium within the meQTL.5a interval are between 114.5–116.5 Mb on Chr5 (**Data S2**). This is the location of genes such as the hepatic TF *Hnf1a*, the sirtuin gene *Sirt4*, heat shock protein *Hspb8*, and the coenzyme *Coq5* (**Fig 1g**). For candidate gene ranking, we narrowed to the 114.5–116.5 Mb peak interval and retained positional candidate genes that (1) have missense or protein truncating variants that segregate in the BXDs, and/or (2) are modulated in expression by a cis-eQTL. This identified 20 positional candidates located in the peak region within meQTL.5a (**Table 2**). 14 of these had missense mutations, and we further used the SIFT (Sorting Intolerant From Tolerant) score to predict the potential deleterious effects on protein function.^57^ SIFT scores range from 0 to 1 with low values (<0.05) predicted to be deleterious. Variants with low SIFT scores are in *Oasl2*, *Srsf9*, *Pxn*, *Rab35*, *Cit*, and *Prkab1*, and the variant in *Hnf1a* also had a comparatively low SIFT score (**Table 2**).

**Table 2.**
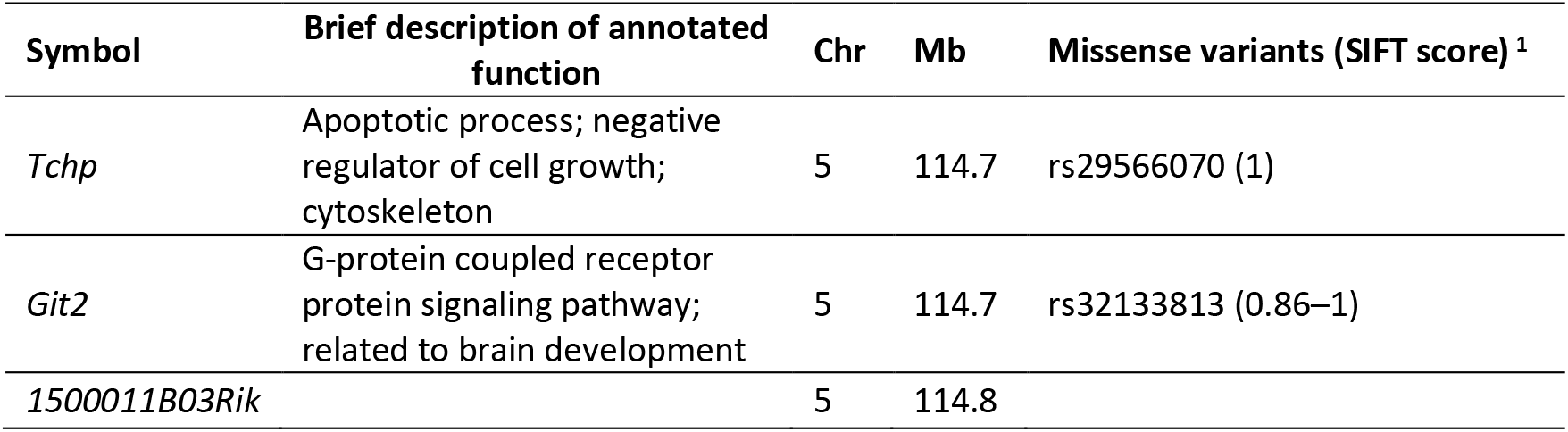
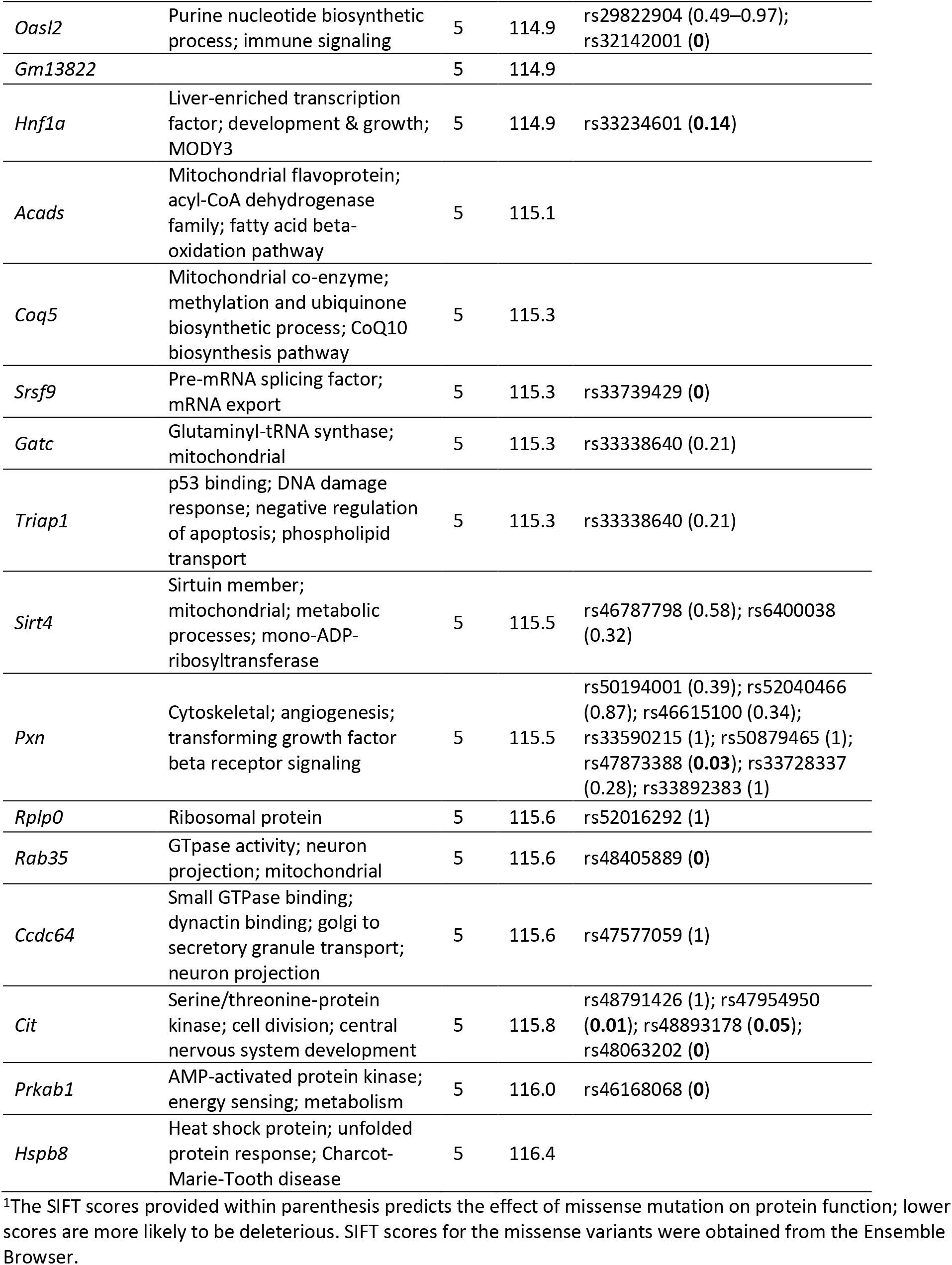
Candidate genes in meQTL.5a

Our next goal was to determine which of these candidate genes formed the most cohesive functional network(s) with the trans-modulated CpGs. For this, we took the list of genes cognate to the trans-CpGs (i.e., gene in which the CpG in located, or the nearest gene if CpG is intergenic), and the list of positional candidates, and searched the STRING database for protein-protein interactions (PPI).^58^ This resulted in a large and highly connected network with an average node degree of 4.21 (PPI enrichment p < 1e-16), and a high enrichment in developmental genes. The central hub was around the trans-modulated *Crebbp*, which had the highest degree of nodes at 47 (**Fig S3**). In this CpG-based network, the candidate gene with the highest degree of connections was *Hnf1a* (13 nodes; **Table S5**). The most enriched KEGG pathway was ‘maturity onset diabetes of the young’ or MODY (mmu04950), and 6 members in the central hub were members of this pathway (**Fig 4a**). This included the positional candidate, HNF1A, which is the causal gene for MODY3.^44^ Enriched GO terms included metabolic processes, cell differentiation, and developmental processes (**Data S9**). *PXN* was another candidate gene in meQTL.5a with high connectivity, but it formed a more peripheral cluster (**Fig S3**). Other candidates such as *Sirt4* and *Hsbp8* had 2 and 0 connections, respectively (**Table S5**). A CpG located in an exon of the hub gene, *Crebbp* (cg27201505), mapped as a strong trans-meQTL to meQTL.5a; however, the *Crebbp* transcript had low expression in adult liver, and the mRNA had only a weak eQTL in meQTL.5a (–Log10p = 2.09 at the meQTL.5a peak interval) (**Fig 4b**). Similarly, a CpG (cg12712768) located in the promoter of *Hnf1a* mapped as a strong cis-meQTL, but the *Hnf1a* mRNA only had weak evidence of cis-modulation (**Fig 4c**). The cis-modulated CpG in *Hnf1a* had a strong positive correlation with the trans-modulated CpG in *Crebbp* (**Fig 4d**), and there was also strong positive correlation between their transcripts (**Fig 4e**). However, the inter-omic correlations between CpGs and transcripts were relatively modest, and although CREBBP formed the central node in the PPI network, the expression of *Crebbp* was uncorrelated with its cognate CpG, and instead, the *Crebbp* transcript had a modestly significant inverse correlation with the *Hnf1a* CpG (**Fig 4f**). The *Hnf1a* transcript was also modestly correlated with its CpGs (**Fig 4g**).

**Fig 4.**
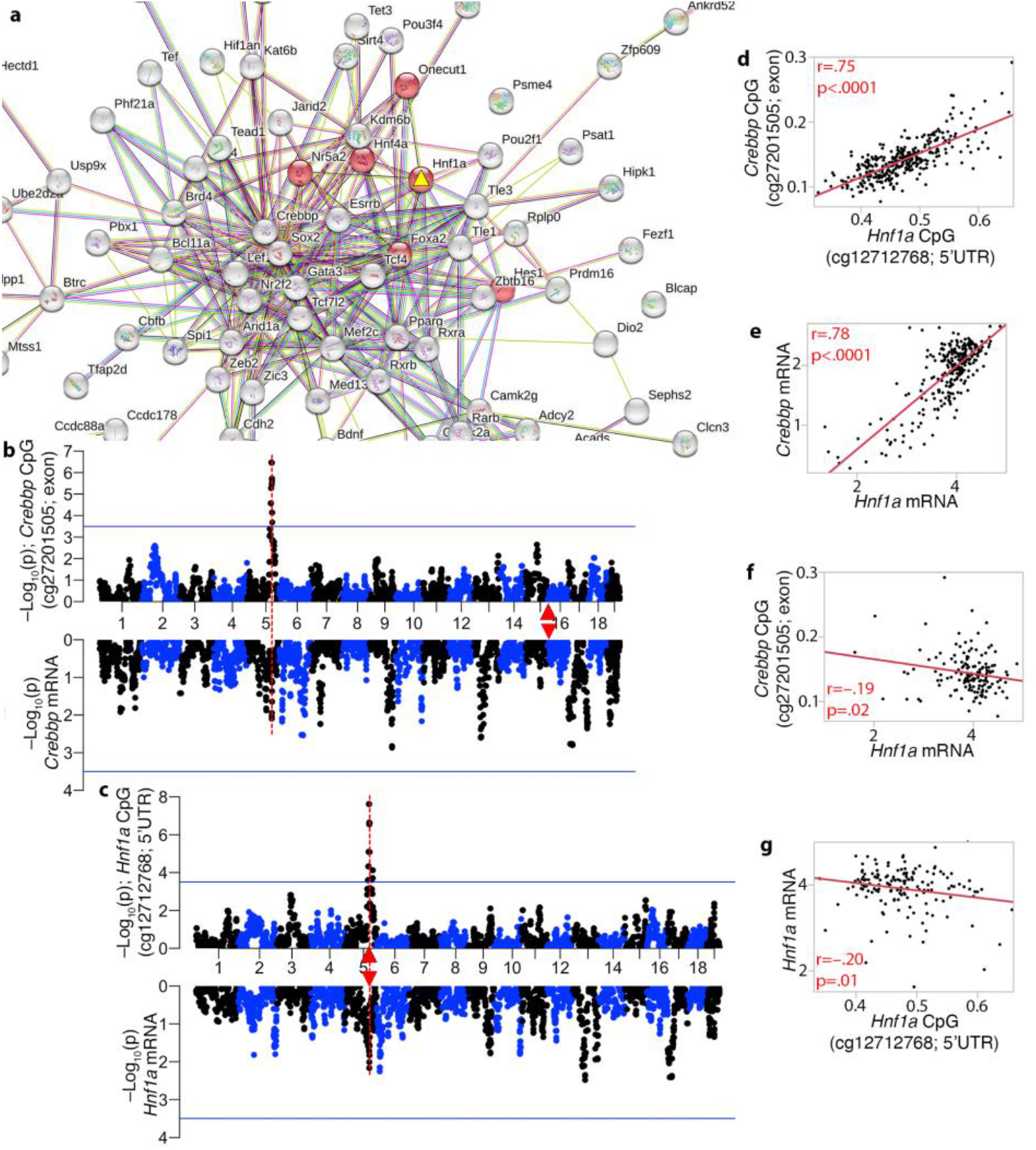
Interaction networks that connect the trans-meQTLs to the meQTL.5a candidates. **(a)** The trans-modulated CpGs and candidate genes in meQTL.5a form a highly connected and functionally enriched protein-protein interaction network. HNF1A (yellow triangle) is the most connected candidate in the central sub-network, and member of the enriched MODY (Maturity Onset Diabetes of the Young) pathway (red nodes). **(b)** CREBBP is the hub gene in the meQTL-based network, and a CpG in its exon is trans-modulated by meQTL.5a (top). Its transcript (bottom of mirrored Manhattan plot) has a weak peak in meQTL.5a. *Crebbp* is located on chromosome 14 (red triangle). **(c)** The CpG in the 5’UTR of *Hnf1a* is cis-modulated, and its expression has a weak cis-eQTL. Strong positive correlation between the CpGs **(d)**, and between the mRNAs **(e)** of *Hnf1a* and *Crebbp*. Weak inverse correlation between **(f)** the *Hnf1a* mRNA and *Crebbp* methylation, and **(g)** between the mRNA and methylation of *Hnf1a*.

We performed a similar PPI analysis for the lists of genes with trans-eQTLs, and trans-pQTLs in meQTL.5a. The trans-modulated mRNAs formed a network with an average node degree of 2.42 (PPI enrichment = 1e-06; **Fig S3**). The trans-modulated proteins formed a smaller but highly connected network with average node degree of 2.34 (PPI enrichment = 5.2e-10; **Fig S4**). At both the transcriptomic and proteomic levels, HNF1A no longer occupied a central position (only one degree of connection for HNF1A in both), and instead, OASL2 was the most connected positional candidate for networks defined from the trans-eQTL and trans-pQTL (**Fig S3, Fig S4**). The functional profiles of the networks were also altered and the most enriched KEGG pathway in the eQTL-based PPI network was autophagy, and the pQTL-based PPI network was enriched in metabolic pathways (**Data S8**). The eQTL and pQTL networks shared similarities; for instance, there was a clique of proteasome subunits (PSMB8, PSMB9, PSMB10) connected to OASL2, and suggests overlapping interactional and regulatory networks at the transcriptomic and proteomic levels that are disconnected from the developmental networks at the methylome level. Overall, this suggests that *Hnf1a* is a strong positional candidate for the trans-meQTLs, but not for the pathways that connect the trans-modulated expression traits.

Due to the apparent centrality of HNF1A within the CpG network, we searched the STRING database for the top 10 high-scoring interaction partners for HNF1A (**Fig 5a**). The present array targets only a few highly conserved CpGs in each of these genes. But even with this sparse profiling of CpGs, 6 of the top 10 PPI interaction partners of HNF1A mapped as trans-meQTLs to meQTL.5a (two members, PCPD1 and PPARA, did not have meQTL data as no CpG probes in the mammalian array targeted these genes). We performed pair-wise expression correlations for these 10 interaction partners and *Hnf1a* using the liver RNA-seq data, and the transcripts formed a highly interconnected network in which the mRNA for *Hnf1a* was connected to 9 of the 10 PPI-based members at |r| ≥ 0.5 (**Fig 5b**). As was the case for the *Crebbp* and *Hnf1a* transcripts, the mRNAs of *Foxa2* (**Fig 5c**) and *Hnf4a* also mapped as weak trans-eQTLs to the same interval (GEMMA based linkage statistics in **Data S6**). For *Gata4*, its mRNA had a relatively strong trans-eQTL in meQTL.5a (**Fig 5d**).

**Fig 5.**
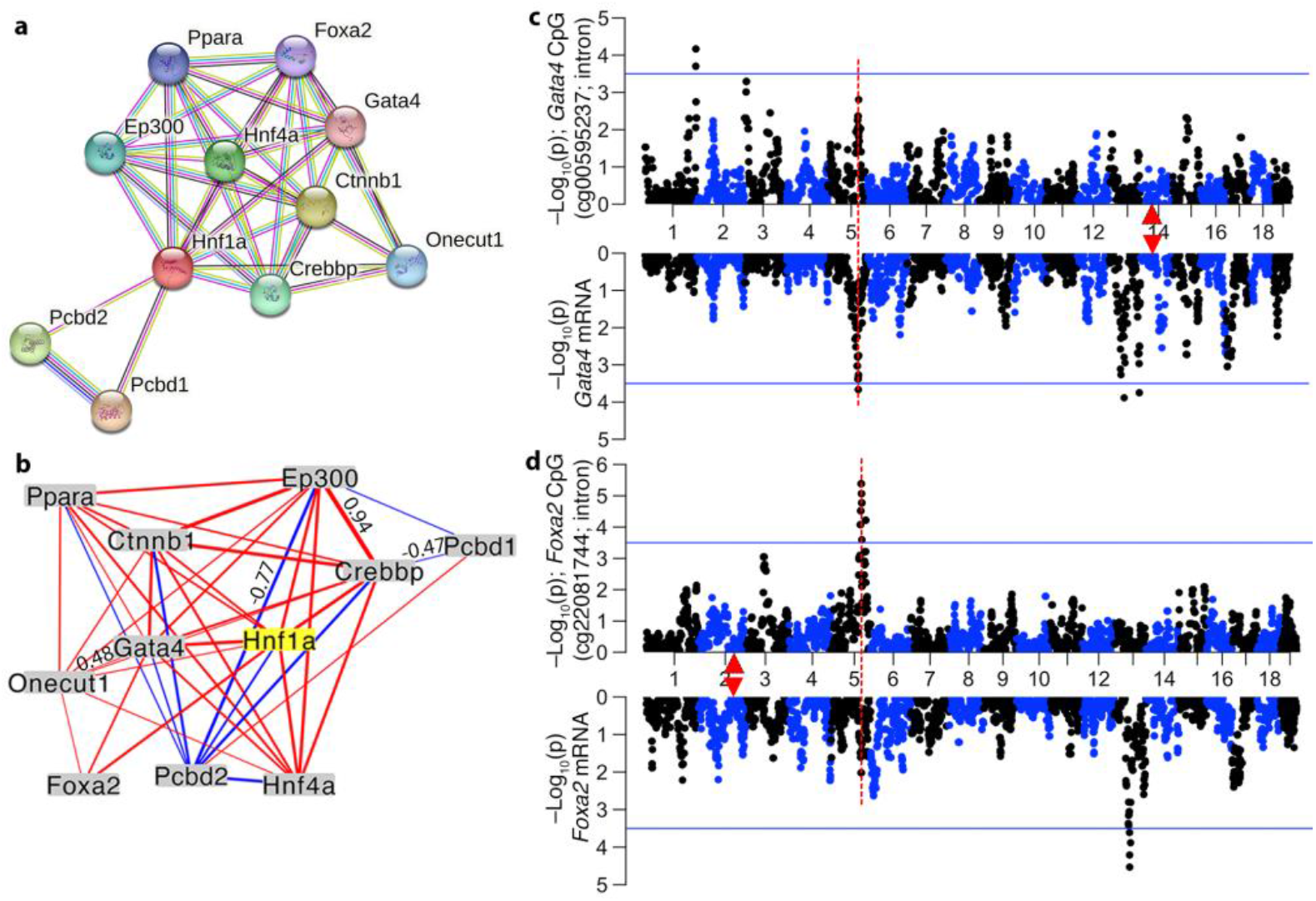
Primary protein-protein interaction partners of HNF1A and their trans-modulation. **(a)** The network shows the top 10 interaction partners of HNF1A based on protein-protein interactions. CpGs in CREBBP, FOXA2 (HNF3B), GATA4, HNF4A, ONECUT1, and PCDB2 are trans-modulated by the meQTL.5a locus, making *Hnf1a* a prime candidate. **(b)** At the transcriptomic level, expression of these genes in the liver are also highly intercorrelated. Only correlations |r| > 0.45 are shown (line thickness conveys strength of correlation; blue: negative; red: positive). Mirrored meQTL (top), and eQTL (bottom) for two example gene: **(c)** *Gata4*, and **(d)** *Foxa2*. Red triangles mark the location of genes.

While we cannot dismiss the other genes highlighted in **Table 2**, *Hnf1a* stands out as a strong candidate for the trans-meQTLs. Our observations suggest that meQTL.5a modulates the methylation, and to a lesser extent, the expression of genes that functionally interact with HNF1A. The missense mutation in *Hnf1a* (rs33234601) results in a proline to serine substitution, with proline as the conserved amino acid across most mammals (based on the comparative genomics track on the UCSC Genome Browser).^59^

### Interaction with aging, diet, and potential impact on longevity

We next examined how the trans-CpGs interface with aging and diet, and how these may potentially influence physiological traits and lifespan. Based on the level of overlap with DMCs that we have previously defined,^6^ the CpGs trans-modulated by meQTL.5a had 3-fold higher enrichment in age-gain CpGs (hypergeometric p = 5.4e-92). There were also significant enrichments in weight- and diet-CpGs, and modest enrichment in age-loss CpGs, but no DMCs related to strain dependent life expectancy (**Fig 6a; Table S6**). Intriguingly, for the age dependent trans-CpGs, whether a site gained or lost methylation with aging depended on the allele effect of meQTL.5a (**Fig 6b**). Trans-CpGs with *D* positive additive effect were more likely to gain methylation with age, whereas the few trans-CpGs with higher methylation for the *B* allele were associated with decrease in methylation with aging (**Fig 6b; Data S2**). This is not due to spurious co-segregation between genotype in meQTL.5a since there is not difference in mean age between the samples with the *DD* genotype (421 ± 170 days) and those with the *BB* genotype (425 ± 184) (**Data S1**). This pattern of allele-dependent effect of age is exemplified by the CpG located in the 5’UTR of *Jarid2*, a canonical member of the Polycomb-Repressive Complex 2 (PRC2) (**Fig 6c**).^60^ This CpG gained methylation with age, and across all ages, mice with the *D* allele in meQTL.5a tended to have higher methylation. An meQTL map for the *Jarid2* CpG using GEMMA is displayed in **Fig 6d**. The expression of *Jarid2* in adult liver was low, and showed no covariance with age, but the mRNA mapped as a weak trans-eQTL to meQTL.5a (p = 0.01). We have previously reported that being fed HFD augments the age-dependent gains in methylation such that HFD results in a more aged methylome.^6, 43^ For the meQTL.5a trans-CpGs, all the CpGs associated with higher methylation for the *D* allele were also increased in methylation by HFD (**Fig 6e**). These trans-CpGs with higher methylation for the *D* allele were also more likely to inversely correlate with body weight (**Fig 6f**). Overall, this pattern suggests a genotype dependent susceptibility of CpGs to the effects of aging and diet.

**Fig 6.**
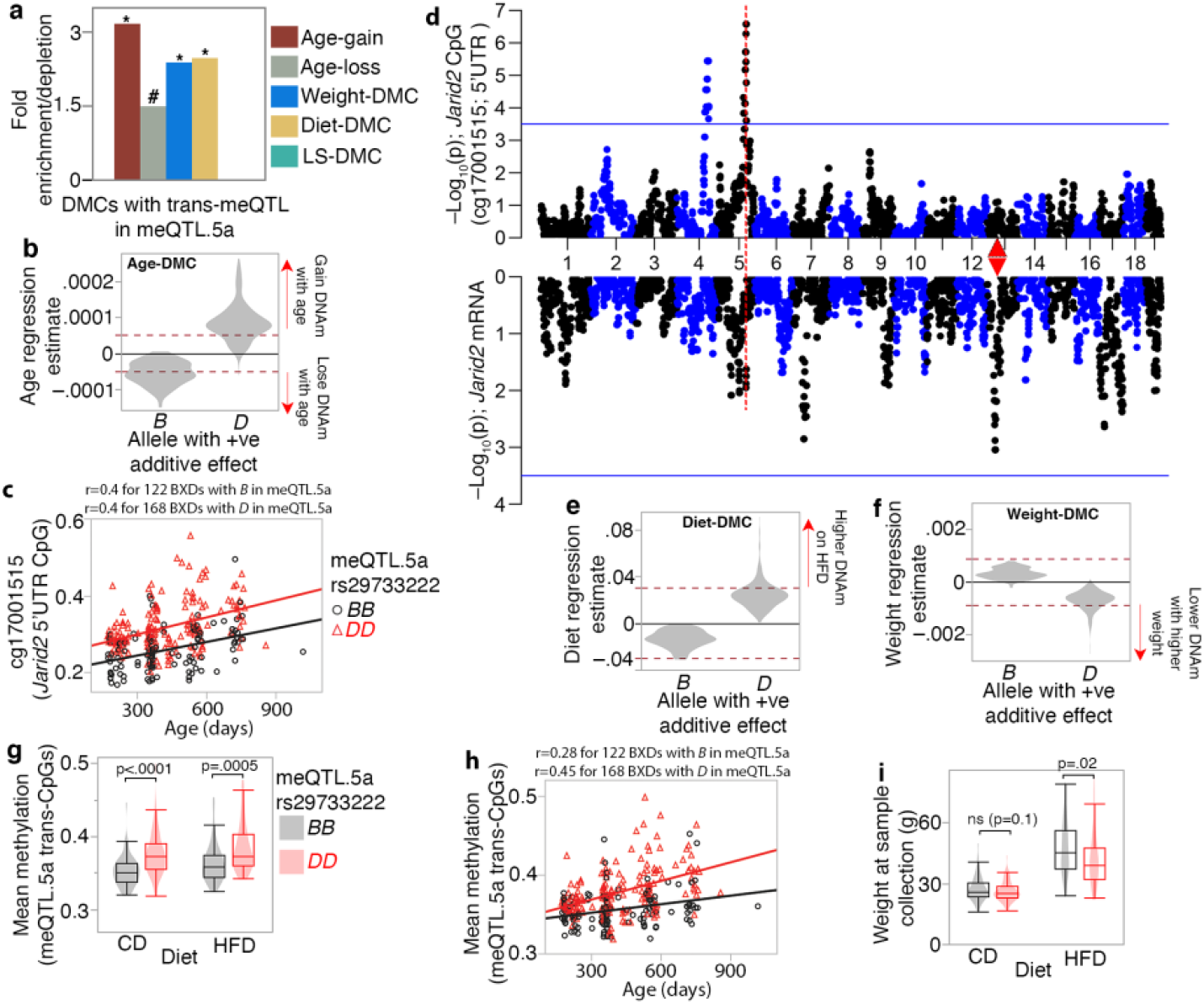
Joint modulation of CpGs by genetic and non-genetic variables. **(a)** CpGs with trans-meQTLs in meQTL.5a are enriched in differentially methylated CpGs (DMCs) associated with aging, diet, and body weight. Asterisks denote hypergeometric enrichment p ≤ 0.001; hash denotes p = 0.006. **(b)** Trans-CpGs that are increased in methylation by the *D* allele in meQTL.5a gain methylation with age (positive regression estimate on the y-axis) while those with higher methylation for the *B* allele lose methylation with age. Dashed red horizontal line indicate Bonferroni p ≤ 0.05 for DMC. **(c)** CpG in the *Jarid2* 5’UTR gains methylation with age and has higher methylation in BXDs with the *DD* genotype in meQTL.5a. **(d)** QTL plots for the *Jarid2* CpG (top Manhattan plot), and mRNA (bottom). Red triangle marks the location of *Jarid2*. **(e)** Allele dependent increase in methylation at the meQTL.5a trans-CpGs due to HFD. **(f)** Allele dependent association with body weight for the trans-CpGs. **(g)** Overall mean methylation of the trans-modulated CpG is higher for BXDs with *DD* genotype in meQTL.5a for both control diet (CD; 0.35 ± 0.02 for *BB*; 0.38 ± 0.03 for *DD*; pair-wise p < 0.0001) and high fat diet (HFD; 0.36 ± 0.03 for *BB*; 0.38 ± 0.03 for *DD* pair-wise p < 0.005). **(h)** Allele dependent increase in mean methylation for the CpGs with trans-meQTL in meQTL.5a. **(i)** Body weight is slightly higher for BXDs with *BB* in meQTL.5a for both CD (27 ± 6 g for *BB*; 26 ± 5 g for *DD*; pair-wise p < 0.1) and HFD (47 ± 13 g for *BB*; 42 ± 13 g for *DD*; pair-wise p < 0.02).

To explore this further, we computed the mean methylation value for all 502 trans-CpGs targeted by meQTL.5a. As expected, the *DD* genotype in meQTL.5a had higher average methylation than the *BB* genotype for both diet groups (**Fig 6g**), and similar to *Jarid2*, the mean methylation increased with age but was overall higher for the *DD* genotype (**Fig 6h**). Body weight was not directly correlated with the mean methylation of the trans-CpGs, despite the enrichment in DMCs inversely correlated with body weight among the trans-CpGs. A multivariable regression showed that the strongest predictor of mean methylation of the trans-CpGs was age, followed by the meQTL.5a genotype, and then diet, but not body weight (**Table 3**). If we treat body weight as the outcome variable, there is only a slightly higher weight for the *BB* genotype (**Fig 6i**), and a multivariable regression showed a modestly significant association between weight and meQTL.5a (**Table 3**). This suggests a pleiotropic effect of meQTL.5a on DNAm and body weight with contrasting allelic effects.

**Table 3.**
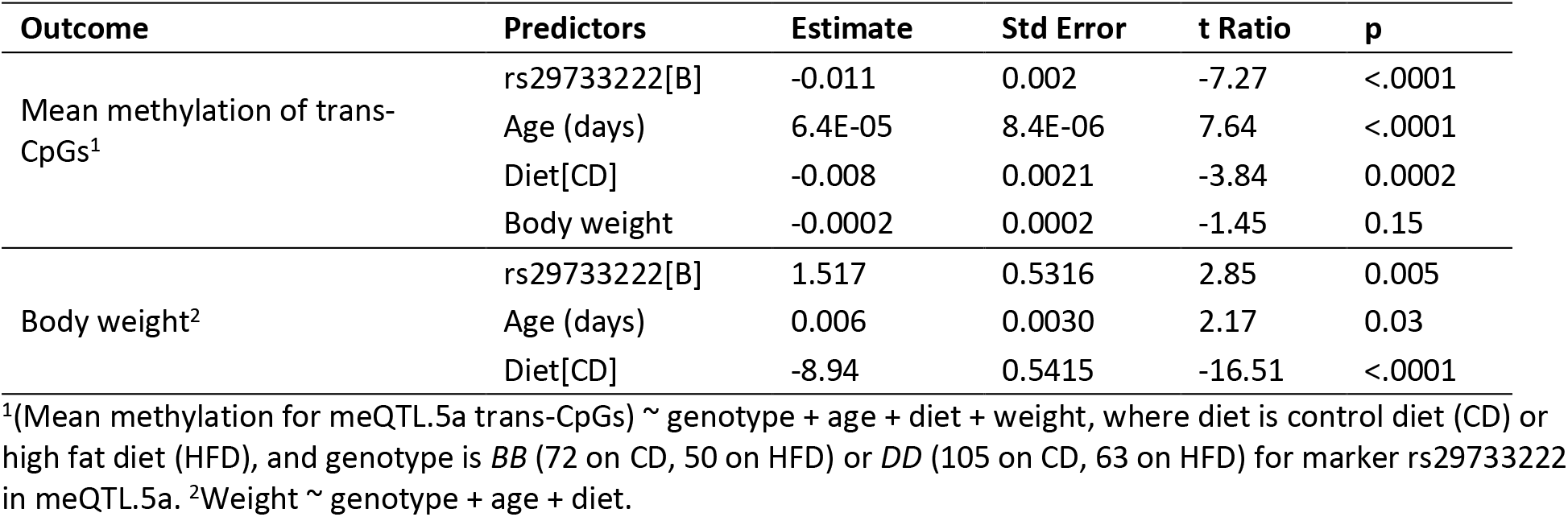
Multivariable variable regressions for mean methylation and weight

| Outcome | Predictors | Estimate | Std Error | t Ratio | p |
| --- | --- | --- | --- | --- | --- |
| Mean methylation of trans-CpGs<sup>1</sup> | rs29733222[B] | -0.011 | 0.002 | -7.27 | <.0001 |
|  | Age (days) | 6.4E-05 | 8.4E-06 | 7.64 | <.0001 |
|  | Diet[CD] | -0.008 | 0.0021 | -3.84 | 0.0002 |
|  | Body weight | -0.0002 | 0.0002 | -1.45 | 0.15 |
| Body weight<sup>2</sup> | rs29733222[B] | 1.517 | 0.5316 | 2.85 | 0.005 |
|  | Age (days) | 0.006 | 0.0030 | 2.17 | 0.03 |
|  | Diet[CD] | -8.94 | 0.5415 | -16.51 | <.0001 |

<sup>1</sup>(Mean methylation for meQTL.5a trans-CpGs) ~ genotype + age + diet + weight, where diet is control diet (CD) or high fat diet (HFD), and genotype is *BB* (72 on CD, 50 on HFD) or *DD* (105 on CD, 63 on HFD) for marker rs29733222 in meQTL.5a. <sup>2</sup>Weight ~ genotype + age + diet.

We obtained bodyweight and lifespan data from a different BXD cohort that were allowed to survive until natural mortality.^35^ We segregated the samples into two groups based on homozygous genotype at the meQTL.5a marker, rs29733222: *BB* (n = 801) or *DD* (n = 972), and tested two predictions: (1) that the *BB* genotype in meQTL.5a will be associated with higher body weight at age 6 months, and (2) although higher body weight is associated with shorter lifespan, we predicted that at this locus, based on a more “aged methylome”, the *DD* genotype will have slightly shorter lifespan. As the longevity cohort has no DNAm data, we could not directly verify whether the *DD* in this group indeed have a more aged methylome, and this is purely an assumption based on the meQTL data. As predicted, the *BB* genotype had significantly higher mean body weight (**Fig 7a, Table 4**). Also consistent with prediction, the *DD* genotype was associated with a slightly higher risk of death between the ages of 650 and 1100 days compared to the *BB* genotype, but only in the CD group (Log-Rank p = 0.008 for CD; Log-Rank p = 0.57 for HFD; **Fig 7b, 7c**). This small difference in the CD group resulted in a median lifespan of 702 days and maximum LS of 1250 days in the *BB* genotype (n = 399), compared to a median of 685 days and maximum of 1197 days in the *DD* genotype (n = 510). Using a multivariable regression model with genotype, diet, and bodyweight at 6 months as predictors, we find that the strongest predictors of lifespan were weight at 6 months, followed by diet, and then by the meQTL.5a genotype (**Table 4**). Given the strong association between body weight at young age and longevity,^35^ the lifespan advantage for the *BB* genotype becomes more apparent when we adjust the longevity data for the weight at 6 months (**Fig 7d**). For the CD mice, after adjusting for weight, the *BB* mice were predicted to have median lifespan that was 46 days longer the *DD* mice (log-rank p < 0.0001). On HFD, the median lifespan of *BB* was only 7 days longer than the *DD* mice (log-rank p < 0.03). Our results suggest a pleiotropic effect of a genetic locus on multiple traits that are also modified by diet and aging. Here we use the terminology by Tyler et al.,^61, 62^ and the model in **Fig 7e** depicts a horizontal pleiotropic effect on body weight and CpG methylation that can moderate the association between this locus and lifespan. We suggest a modest vertical pleiotropic effect on lifespan that is mediated by the methylome, and the genotype with the less aged methylome (*BB*) having a slight lifespan advantage.

**Fig 7.**
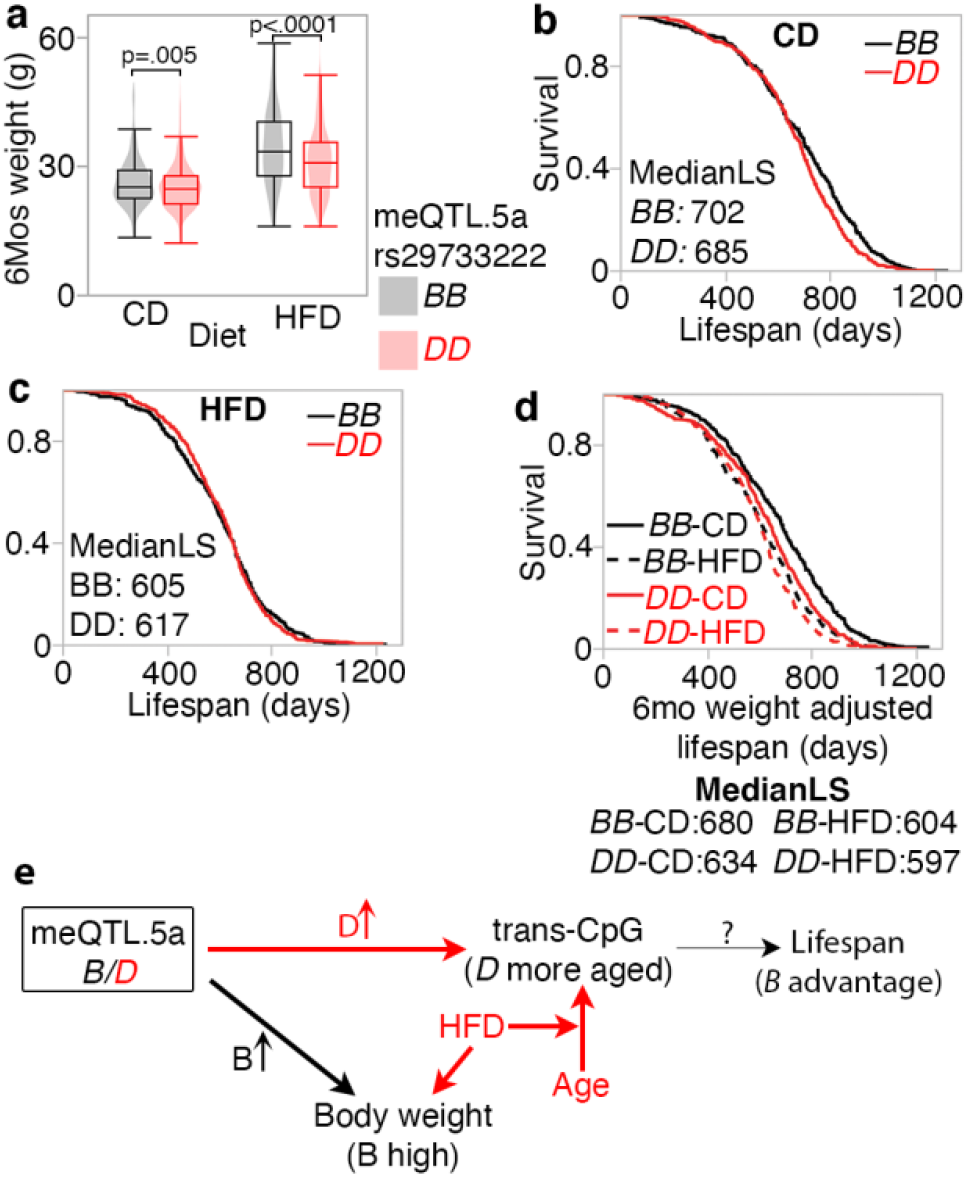
Pleiotropic influence on meQTL.5a on body weight at young age and lifespan. **(a)** Body weight at 6 months (mos) from a separate cohort of BXD mice show higher mean weight for strain with *BB* genotype in meQTL.5a for control diet (CD; 26 ± 6 g for *BB*; 25 ± 5 g for *DD*; pair-wise p < 0.004) and High fat (HFD; 34 ± 9 g for *BB*; 31 ± 8 g for *DD*; pair-wise p < 0.0001). Samples numbers: 383 *BB* and 503 *DD* for CD; 392 *BB* and 457 *DD* for HFD. **(b)** Kaplan-Meir survival plots by genotype at meQTL.5a for CD mice (399 *BB* and 510 *DD*). Median lifespan in days (MedianLS) for the genotypes shown (log-rank p = 0.008). **(c)** Similar survival plot for HFD shows no significant difference between genotypes (402 *BB* and 462 *DD*). **(d)** Kaplan-Meir survival after adjusting lifespan for 6 mos weight. Adjusted median lifespan in days shown for each genotype-by-diet below the graph. Within each diet, the *BB* genotype has longer lifespan compared to *DD* (based on pair-wise comparison, log-rank p < 0.0001 for CD, and p = 0.03 for HFD). **(e)** Model depicting horizontal pleiotropic influence on CpG methylation and weight, and vertical pleiotropic influence on lifespan mediated by CpG, which are also under the influence of aging and diet.

Table 4.

**Table 4.**
| Outcome | Term | Estimate | Std Error | t Ratio | p |
| --- | --- | --- | --- | --- | --- |
| Weight_6M<sup>1</sup> | rs29733222[B] | 0.98 | 0.17 | 5.78 | <.0001 |
|  | Diet[CD] | -3.38 | 0.17 | -20.11 | <.0001 |
| Lifespan<sup>2</sup> | rs29733222[B] | 14 | 4.41 | 3.08 | 0.002 |
|  | Diet[CD] | 26 | 4.82 | 5.43 | <.0001 |
|  | Weight_6M | -4.0 | 0.62 | -6.34 | <.0001 |

<sup>1</sup>(Weight at 6 months) ~ genotype + diet. <sup>2</sup>Lifespan ~ genotype + diet + weight at 6 months, where diet is control diet (CD) or high fat diet (HFD), and genotype is *BB* (399 on CD, 402 on HFD) or *DD* (510 on CD, 462 on HFD) for marker rs29733222 is meQTL.5a.

### Phenome-wide association analysis for *Hnf1a*

The BXDs have accrued a vast collection of traits over decades, and we next performed a phenome-wide association analysis (PheWAS) to identify other higher-order traits that may be modulated by the meQTL.5a interval.^63^ Note that although we used “*Hnf1a*” as the term in the PheWAS search,^64^ the results are from family-based linkage mapping (not allelic associations), and the linkages are to relatively large QTL intervals close to the *Hnf1a* locus, and not to an *Hnf1a* allele. At −Log_10_p ≥ 3, there were 10 BXD traits that included one immune related phenotype, four traits related to the nervous system, and five metabolic traits related to fat content and amino acid ratios (**Table 5**). The strongest QTL was for susceptibility to rickettsia infection;^65^ however, the peak region for this trait was proximal to the meQTL.5a interval (∼104 Mb; GEMMA based linkage statistics is **Data S6**). The brain related traits were also little distant from meQTL.5a (at ∼107 Mb for brain activity measured by Ito et al.,^66^and at 117 Mb for cell proliferation^67^). The metabolic traits peaked at the meQTL interval and these included measures of fat content in liver, and ratio of branched chain amino acids to total amino acids (**Data S5, Fig S6**).^68–70^ Although the QTLs for the metabolic traits are suggestive, it does indicate a potential role for the meQTL.5a region in higher order metabolism, and similar to the higher body weight, the *BB* genotype had higher liver fat content.

**Table 5.** Phenotypes that map to meQTL.5a

| GN number<sup>1</sup> | Phenotype | Category | PMID | -log10(p) | QTL peak location |
| --- | --- | --- | --- | --- | --- |
| 17439 | Rickettsia tsutsugamushi susceptibility of both sexes at 6-12 weeks-of-age | Immune | 6774020 | 4.42 | Peak at 104 @rs32034514 |
| 17266 | Brain activity and coherence of electrical field oscillations at 160 Hz in L2/3 of the primary whisker motor cortex | Nervous system | 24686563 | 4.03 | Peak at ~107 Mb @rs29681689 |
| 15092 | Total fat content measured by Fourier Transform Infrared Spectroscopy in liver at 140 days, males, fed high fat diet fed from 4 weeks on | Metabolism | 23758785 | 3.94 | Peak from 110–114 Mb |
| 15091 | Saturated fat content measured by Fourier Transform Infrared Spectroscopy in liver at 140 days, males, fed high fat diet feeding from 4 weeks on | Metabolism | 23758785 | 3.62 | Peak from 110–114 Mb |
| 15094 | Ratio of lipid to protein content in liver at 140 days, males, fed high fat diet from 4 weeks on | Metabolism | 23758785 | 3.44 | Peak from 110–114 Mb |
| 14786 | Proliferation of BrdU-labeled cells in subgranular zone, 1h BrdU injection, unadjusted data | Nervous system | 24640950 | 3.43 | Peak at 117 Mb @rs29728022 |
| 17265 | Brain activity and coherence of electrical field oscillations at 159 Hz in L2/3 of the primary whisker motor cortex | Nervous system | 24686563 | 3.31 | Peak at ~107 Mb @rs29681689 |
| 16783 | Ratio of total branched-chain amino acid/total amino acid | Metabolites | 22939713; 30709776 | 3.23 | Peak at 110 Mb @rs49420585 |
| 17245 | Brain activity and coherence of electrical field oscillations, coherence at 139 Hz between local field potentials (LFP) at two sites (0.3 mm apart) in L2/3 of the primary whisker motor cortex in awake 6 | Nervous system | 24686563 | 3.19 | Peak at ~107 Mb @rs29681689 |
| 16774 | Ratio of total branched-chain amino acid/Alanine_CD | Metabolites | 22939713; 30709776 | 3.1 | Peak at 110 Mb @rs49420585 |

<sup>1</sup>Search carried out using *Hnf1a* as the search key in https://systems-genetics.org/; the traits ID can be used to retrieve the BXD strain level data from www.genenetwork.org

Complementing the murine family-based PheWAS, we also searched for GWAS hits associated with variants in *HNF1A* using two GWAS databases: GWAS Atlas, and the NHGRI-EBI GWAS Catalog.^71, 72^ At minimum p < 1e-05, the GWAS Atlas identified 85 traits, and the GWAS Catalog identified over 400 traits associated with variants in *HNF1A* and *HNF1A-AS1*, the antisense RNA transcribed from *HNF1A* (**Data S10**, **Data S11**). The trait with the strongest association was C-reactive protein levels, a measure of inflammation and cardiovascular health, followed by levels of Gamma glutamyl transpeptidase (GGT), a measure of liver damage.^73–76^ These were followed by lipid levels and coronary artery disease.^77–79^ Other traits associated with *HNF1A* included age at puberty, birth weight, and diabetes.^80–84^ While not a replicated genome-wide significant hit, a variant near *HNF1A* (rs6489785) is reported to be one of 37 “longevity SNPs” that have a small-effect on human lifespan.^85^ This is consistent with our model where the meQTL.5a locus could make an indirect and modest contribution to lifespan variation.

## Discussion

We have provided an overview of the regulatory loci that influence methylation of conserved CpGs in the murine liver. Overall, the results show complex interrelationships and genetic pleiotropy on DNA methylation and physiological traits. As expected, cis-meQTLs are associated with higher LOD scores compared to the trans-meQTLs.^14^ Given the strong contribution of genotype to the variance of cis-CpGs, the cis-modulated CpGs were depleted in DMCs related to aging and diet. In contrast, the trans-CpGs were enriched in DMCs related to both genetic traits (lifespan, body weight), and non-genetic variables (aging and diet). In terms of genomic location, the cis-CpGs were enriched in intergenic sites, which is consistent with reports from human studies,^8, 15^ and also in intronic regions, and were highly depleted in 5’UTR and bivalent promoter states (PrB), which are regions that are strongly modified by aging and typically show methylation gains over time.^6^ This suggests that aging has a limited impact on CpGs that are under strong cis-modulation. On the other hand, trans-CpGs have multifactorial variation and could present key sites for gene-by-environment interactions.

For meQTL mapping, we implemented a stringent regression model that adjusted for age, diet, weight, and genetic relatedness, and corrected for unmeasured variance by including the top principal component as a cofactor. However, note that compared to the statistical thresholds that are applied in human meQTL studies, we used a rather relaxed threshold of LOD ≥ 3.5 for both cis- and trans-effects. This was because our sample size was modest and our analysis was a family-based linkage mapping done in 41 BXD progeny strains, F1 hybrids, and the parent strains.^33^ If we increase the stringency for the trans-meQTLs to LOD ≥ 4.5, then only 309 strong trans-effects remained and 76 of these (i.e., nearly 25%) were in meQTL.5a. The relaxed statistical threshold is a caveat to keep in mind. For additional evidence, we evaluated the trans-meQTLs for biological coherence and overlap with eQTLs. For instance, many of the immediate interaction partners of HNF1A have weak trans-eQTLs overlapping the trans-meQTLs. The other strategy we used was to reduce the dimensionality of the methylome data by performing an unsupervised clustering of the CpGs into modules and treating the module eigengenes as the representative quantitative traits, and this too supported meQTL.5a as a modulator of functionally connected CpG networks.

Among the meQTL hotspots Eaa19 is also noteworthy. Eaa19 is linked to epigenetic clock acceleration, and potentially influences susceptibility to entropy accumulation in the liver methylome.^6^ In our 2022 paper, we identified several candidate genes in Eaa19. However, like meQTL.5a, Eaa19 is gene dense and harbors several positional candidates. In the present work, we focused on meQTL.5a, and used several strategies to prioritize the most plausible candidates. Without dismissing the other candidates (e.g., *Oasl2*, *Srsf9*, *Pxn*, which contain variants predicted to be deleterious), our analysis led us to *Hnf1a* as a functionally highly relevant prime suspect in meQTL.5a.

### Developmental genes, the methylome, and aging

HNF1A is a member of the hepatocyte nuclear factor family of TFs, and is mainly expressed in the liver, kidney, and pancreas.^86^ Other HNF members include HNF4A, FOXA2 (aka, HNF3B), and ONECUT1, which all have trans-meQTLs in meQTL.5a. During embryonic development, the HNFs and GATA TFs participate in complex autoregulatory networks that modulate the spatial and temporal expression of downstream genes.^23, 86, 87^ Targets of HNF1A include the metabolic and longevity gene, *Igf1* (insulin-like growth factor 1).^87, 88^ MODY3, which is caused by mutations in *HNF1A*, is the most common form of maturity onset diabetes of the young.^89, 90^ *HNF1A* mutations also lead to dysregulation in fatty acid synthesis and transport that can cause fatty acid accumulation in the liver.^86^ A GWAS study also found that a variant in *HNF1A* (rs6489785) is one of 169 variants that jointly contribute to human longevity.^85^ Some mutations in *HNF1A* do not cause MODY but increase the susceptibility to type 2 diabetes and lower BMI.^91, 92^ In mice, deletion of *Hnf1a* causes Laron dwarfism and hyperglycemia.^93–95^

Although HNF1A is not a direct regulator of DNAm, there is some intriguing evidence that it contributes to the epigenetic state. For instance, deletion of *Hnf1a* in mice causes a change in the local chromatin structure and affects the spatial location of its target regions in the nucleus.^96^ Furthermore, a study from 2008 showed that CpGs located in HNF1A binding motifs were hypomethylated in the liver and had tissue-dependent differential methylation that correlated with gene expression.^97^ This suggests that the binding affinity of HFN1A at these sites could influence CpG methylation. Generally, binding of protein factors (e.g., GATA6, CTCF, REST) to motifs that contain CpGs result in low methylation.^20, 23^ In the case of the BXDs, the D2 allele in rs33234601 (Pro423Ser) is the unusual variant as almost all vertebrate species have a proline at this amino acid position, and only few have serine (e.g., squirrel, elephant; based on the Vertebrate Multiz Alignment track in the UCSC Genome Browser).^59^ Expression of *Hnf1a* has a modest cis-eQTL that is associated with positive additive effect for the *B* allele at meQTL.5a.

Many of these CpGs that are trans-modulated by meQTL.5a are characterized by a low methylation profile (“hypomethylated” with methylation beta-scores closer to 0), and increase in methylation with aging (illustrated by the *Jarid2* CpG, and the mean methylation of the trans-CpGs in **Fig 6**). Since binding by TFs generally result in lower methylation at the biding motifs,^20, 23^ we could speculate that the *D* variant of HNF1A has a lower DNA binding affinity, and the BXD strains with *DD* at meQTL.5a could begin life with heightened methylation at the target sites. If we consider this in terms of epigenetic entropy, then a hypomethylated state presents a low entropy landscape.^6^ For the *DD* genotype however, the methylation beta-values at these CpGs will be closer to 0.5, and will approach a more random epigenetic state at an earlier age compared to strains that have a *BB* genotype at meQTL.5a.

An interesting feature of the *Hnf1a* gene is that the promoter and first intron overlaps the long non-coding RNAs (lncRNA), *Hnf1aos1* and *Hnf1aos2*.^98^ The cis-regulated CpG in *Hnf1a* (shown in **Fig 4c**) is in this lncRNA, and the RNA products have been shown to have a cis-acting regulatory role and implicated in cell proliferation and tumor progression.^98, 99^ Furthermore, *Hnf1aos1* interacts with EZH2, the catalytic subunit of PRC2, in liver tissue.^100^ Genes that are regulated by PRC2, and CpG sites that interact with EZH2 are known to be highly susceptible to age-dependent increases in methylation,^101–103^ and the lncRNA is another plausible link between *Hnf1a* and the epigenome. Notably, one of the strongest trans-modulated CpGs is located in *Jarid2*, a member of the PRC2 complex,^60^ and we can see that while the CpG in *Jarid2* gains methylation with age, the *DD* strains start out with a higher methylation compared to the *BB* strains (see **Fig 6c**). This presents links between a development TF and the PRC2 complex that suggests deeper connections between epigenesis (i.e., embryonic development), and the aging of the epigenome.

### Pleiotropy on CpGs and physiological traits

In the BXDs, low body weight at young age predicts longer lifespan and slower epigenetic aging.^6, 35, 104^ However, the meQTL.5a interval has contrasting allelic effects on body weight and lifespan. Specifically, despite the higher body weight and higher liver lipid levels for the *B* allele in meQTL.5a, it is the *D* allele that is associated with slightly shorter lifespan. These effect on weight (specifically, lower body weight) is also seen in *Hnf1a*-null mice, and *HNF1A* variants in humans. Generally, when downstream targets of *Hnf1a* are deleted, it results in smaller stature and longer lifespan in both humans and mice. For instance, deficiency in growth hormone or IGF1 confers longer lifespan and healthspan.^105, 106^ In some instances of Laron syndrome (LS), individuals exhibit insulin resistance and hyperlipidemia but still have long lives.^107, 108^ Mouse models of Laron dwarfism also age slower and have longer lifespan.^109, 110^ However, direct deletion of *Hnf1a* in mice, or deleterious mutations in *HNF1A* in humans, do not appear to confer any lifespan advantage despite the mice having a form of Laron dwarfism and humans have lower BMI.^91–95^

We suggest that HNF1A, in addition to its role as a TF for developmental and metabolic genes, also influences the epigenome early in life, and contributes to epigenomic maintenance in adulthood and aging. We present a model in which the meQTL.5a locus exerts horizontal pleiotropic effects on physiological traits and CpG methylation. The pleiotropic influence results in the *D* allele increasing methylation at sites that typically have low methylation when young, and the *B* allele increasing body weight and lipid levels. The methylation of the target CpGs, which are also under convergent influence of aging and diet, then contribute to variation in survival trait, with the *D* allele associated with slightly shorter lifespan. In this model, lifespan is a distal complex trait that shows only a modest linkage to meQTL.5a, while the intermediate traits (the CpGs) have a stronger linkage.

In conclusion, we have identified meQTL.5a as a trans-meQTL hotspot that modulates several CpGs in trans. The pleiotropic effect of meQTL.5a could contribute to variation in body size, metabolic traits, CpG methylation and lifespan. *Hnf1a* is a key candidate in this locus, and the potential influence of the HNFs on the epigenomic state during development could contribute to aging and longevity.

## Methods

### Description of DNAm samples and data

The data we use in this study has been previously reported, and the full data is available from NCBI Gene Expression Omnibus (GEO accession ID GSE199979).^6^ In brief, these are liver DNAm data generated on the HorvathMammalMethylChip40 array from 339 mice that belong the BXD Family. Information on each animal (strain, age, weight, diet, etc.) along with all relevant variables used in this study are provided in **Data S1.**

### Methylation QTL mapping with R/QTL2

Each CpG was mapped against 7127 informative autosomal genotype markers distributed across the autosomal chromosomes using the R/qtl2 software.^45^ The full methylation data is available from the NCBI Gene Expression Omnibus database (GEO accession ID GSE199979), and the genotype data used from mapping is provided as **Data S12**. We performed QTL mapping using a univariate linear mixed model that accounts for genetic relatedness. We first computed genotype probabilities and employed that to obtain genetic relatedness matrices (GRM), or the kinship, using a Leave One Chromosome Out (LOCO) scheme. Genome scans included age, diet and the top PC as covariates (variables provided in **Data S1**), and were implemented using a ‘scan1’ function with genotype probabilities as input while adjusting for relatedness outside the chromosome of interest. We next estimated genetic effects and genetic directions between the two genotypes was computed as (*DD* – *BB*). The R codes are provided as Supplemental information (**Data S13**).

### CpG co-methylation networks

We used the WGCNA R package to cluster the CpGs into inter-correlated modules.^46^ The full set of CpGs (∼28K) was used for network definition. Prior to WGCNA, we performed hierarchical clustering (hclust function in R with method = “average”) for outlier detection and excluded one sample (UT153). WGCNA first constructs a pair-wise correlation matrix, and this was converted to a scale free adjacency matrix using default parameters, and with a soft power threshold, β = 6. The β = 6 was associated with a mean connectivity of 168, and maximum connectivity of 1560. The adjacency matrix was converted to a topological overlap matrix (TOM), and the dissimilarity matrix (1 – TOM), and the hclust() function with the “average” method was used to cluster the CpGs. To group the CpGs into modules, we applied the dynamic tree cutting method (cutreeDynamic), with minimum module size = 35, and deepSplit = 2. This resulted in the 14 CpG families (aka, modules), and the grey module, which had 1284 CpGs that did not fit into the other modules. The top principal component was derived from each module and taken as the representative ME. The R codes used are provided as supplementary information (**Data S14**).

### QTL mapping using the Genenetwork web tool

Aside from the main meQTL mapping that was done using R/qtl2, addition QTL analyses were done on the web platform, GeneNetwork, which provide interface to few different mapping algorithms.^34, 52^ We used the GEMMA algorithm, which adjusts for the BXD kinship structure using linear mixed modeling.^50, 51^ The MEs from the WGCA were uploaded to Genenetwork, and QTL mapping for each ME was done with age, diet and body weight (weight at time of tissue collection) as cofactors. Instructions on how to retrieve the ME traits on GeneNetwork are provided in **Data S6**. QTL mapping for the higher order traits identified by the PheWAS was also done using GEMMA, and for these, the data are at the strain levels (i.e., strain means), and instruction on trait retrieval are provided in **Data S6**.

### Enrichment analysis and other statistics

As previously described, we have annotated each CpG by genomic context (i.e., intergenic, 3’UTR, intron, exon, 5’UTR) and chromatin state.^6, 55, 56^ For enrichment analysis, we compared the frequency of these features among the cis- and trans-modulated CpGs relative to the array background (i.e., ∼28K CpGs), and enrichment or depletion p-values were calculated using a hypergeometric test (formulae provided under **Table S1**). In addition to genomic locations, the CpGs have been classified into differentially methylated by age, diet (high fat vs normal lab chow), and body weight based on a multivariable epigenome-wide association analysis.^6^ The frequency of these differentially methylated CpGs among the cis- and trans-modulated CpGs were also compared against the array background using the hypergeometric test (the R codes are provided under **Table S2**). All other statistical tests (Pearson correlations, linear regression modeling, and survival analyses) were done using JMP (version 16).

### Bioinformatic resources

For the meQTL.5a trans-modulated CpGs, and CpGs in the Blue module, biological functions and transcription factor motif enrichment analysis was done using the R package for Genomic Regions Enrichment of Annotations Tool (rGREAT; version 3).^53, 54^ The base coordinate for each CpG was provided (GRCm38/mm10 reference genome), and comparison was against the array background. For the trans-modulated mRNAs and proteins, we used the gene symbol as the identifier, and enrichment analysis was done on DAVID.^111^ Another enrichment analysis to connect the trans-modulated genes with the positional candidates was based on protein-protein networks, and for this, a non-redundant list of the trans-modulated genes and candidate genes was uploaded to the STRING (version 11.5).^112, 113^

For candidate gene selection, we search for cis-eQTL in the BXD liver RNA-seq data using the GeneNetwork search tool.^41^ To identify protein truncating and missense variants located in the positional candidate genes, we use the Ensemble Variant Table tool for the mouse gene (GRCm38) and the Mouse Genome Informatics variant database, and selected the genes that had such variants between B6 and D2.^114–117^ We used the integrated systems genetics web platform to perform a PheWAS for the *Hnf1a* locus in the BXDs.^63, 64^ For human PheWAS, we used *HNF1A* as the search term and retrieved GWAS hits from two databases: the GWAS Atlas, and the GWAS Catalog.^71, 72^

## Supporting information

All supplemental files

## Acknowledgements

The present study used data generated from the biospecimen resource created by the UTHSC BXD Aging Colony (PI: R Williams), and we have benefitted greatly from Dr. Robert Williams’ vision, leadership, and enthusiasm. We thank all members of the team, especially Dr. Lu Lu, Casey Chapman, and Dr. Suheeta Roy. We thank all members of the GeneNetwork Bioinformatics team. This work was funded by the NIA NIH grant R21AG055841. The BXD Aging Colony was funded by the NIA NIH grant R01AG043930.

## Competing Interest

Steve Horvath is a founder of the non-profit Epigenetic Clock Development Foundation, which plans to license several patents from his employer University of California Regents. These patents list SH as an inventor. The other authors declare no conflicts of interest

