## Supplementary material for "Pleiotropic Influence of DNA Methylation QTLs on physiological and aging traits": All supplemental files: DataS13_BXD_DNAm_rqtl2.R

```
library(qtl2)
library(tictoc)
library(foreach)
library(doParallel)


# current working directory: `Analysis`
# data are stored in a 'Data' folder

# read data
bxd<-read_cross2("../Data/dnamnia.yaml")

# check NA's in genotypes: no NA's found
cnames<- chr_names(bxd)
for (i in 1:length(cnames)){
  txt=paste('sum(is.na(bxd$geno$`',cnames[i],'`))')
  print(eval(parse(text=txt)))
}


# compute genotype probabilities 
pr <- calc_genoprob(bxd, error_prob=0.002, cores=5)

# compute kinship by loco excluding X chromosome
k_loco<-calc_kinship(pr,"loco",omit_x=TRUE,cores = 5)

# change the type of each covariate
age<-as.numeric(bxd$covar$`Age (days)`)
diet<-(bxd$covar$Diet == "HFD")*1
pc1<-as.numeric(bxd$covar$PC1)
covar=cbind(age,diet,pc1)
row.names(covar)<- row.names(bxd$covar)
covar=data.frame(covar)


# genome scan adjusting age & diet
tic()
out_ad<-scan1(pr,bxd$pheno ,kinship = k_loco,addcovar = covar[c("age","diet")],cores = 4)
toc()
write.csv(out_ad,file = "bxd_age+diet_loco.csv",sep = ",")

# genome scan adjusting all covariates (age, diet & pc1 )
tic()
out1<-scan1(pr,bxd$pheno,kinship = k_loco,addcovar = covar  ,cores = 10)
toc()
write.csv(out1,file = "bxd_allcovar_loco.csv",sep = ",")


# compute directions of effects for age+diet (on linux) : parallel computing

#create the cluster & register
myCluster <- parallel::makeCluster(12, type = "FORK",outfile="")
registerDoParallel(myCluster)
#check cluster definition (optional)
print(myCluster)

cnames<-chr_names(bxd)[1:19]
pnames<-colnames(bxd$pheno)
end<-ncol(bxd$pheno)

# three for-loops implemented for both genome scans: j=1:10000, 10001:20000, 20001:end
tic()
output0<-foreach(j=20001:end,
                 .combine='rbind')%dopar% {
                   edir<-NULL
                   for (l in 1:length(cnames)) {
                     coef1<-scan1coef(pr[,toString(l)],bxd$pheno[,j],kinship=k_loco[[toString(l)]],addcovar = covar[c("age","diet")])
                     #get directions for BB & DD
                     edir<-c(edir,sign(coef1[,2]-coef1[,1])) 
                   }
                   edir*out_ad[1:7127,j]
                 }
toc()

stopCluster(myCluster)
row.names(output0)<-pnames[20001:end] 

write.csv(output0,file = "signedLODs_age+diet_loco1.csv",sep = ",") # j=1:10000 in for-loop
write.csv(output0,file = "signedLODs_age+diet_loco2.csv",sep = ",") # j=10001:20000
write.csv(output0,file = "signedLODs_age+diet_loco3.csv",sep = ",") # j=20001:end


#compute directions of effects for all covariates (on macbook)

#create the cluster & register
myCluster <- parallel::makeCluster(4, type = "FORK",outfile="")
registerDoParallel(myCluster)
#check cluster definition (optional)
print(myCluster)

cnames<-chr_names(bxd)[1:19]
pnames<-colnames(bxd$pheno)
end<-ncol(bxd$pheno)
# 7127 markers
tic()
tem<-foreach(j=1:10000,
             .combine='rbind')%dopar% {
               edir<-NULL
               for (l in 1:length(cnames)) {
                 coef1<-scan1coef(pr[,toString(l)],bxd$pheno[,j],kinship=k_loco[[toString(l)]],addcovar = covar)
                 #get direction for BB & DD
                 edir<-c(edir,sign(coef1[,2]-coef1[,1]))
               }
               edir*out1[1:7127,j]
             }
toc()

stopCluster(myCluster)
row.names(tem)<-pnames[1:10000]

write.csv(tem,file = "signedLODs_allcovar_loco1.csv",sep = ",") # j=1:10000 in for-loop
write.csv(tem,file = "signedLODs_allcovar_loco2.csv",sep = ",") # j=10001:20000
write.csv(tem,file = "signedLODs_allcovar_loco3.csv",sep = ",") # j=20001:end
```
