## Supplementary material for "Pleiotropic Influence of DNA Methylation QTLs on physiological and aging traits": All supplemental files: DataS14_WGCNA_DNAm.R

```
library(WGCNA)

# read DNA methylation data
d <- data.frame(read.csv(".../27966probes_sesame.csv", header=T))

# subsetting columns with methylation beta-values
dat <- d[,3:341]
row.names(dat) <- d$Cgid

options(stringsAsFactors=FALSE)

# transpose the data, and check
datExpr0 <- data.frame(t(dat))
dim(datExpr0)
datExpr0[1:10, 1:5]
datExpr0[330:339, 1:5]

# check for missing values
gsg = goodSamplesGenes(datExpr0, verbose = 3)
gsg$allOK

# using hierarchical clustering to detect outliers
sampleTree = hclust(dist(datExpr0), method = "average")
jpeg("DNAmLiver_SampleTree.jpg")
par(cex = 0.6)
par(mar = c(0,4,2,0))
plot(sampleTree, main = "Sample clustering to detect outliers", sub="", xlab="", cex.lab = 1.5, cex.axis = 1.5, cex.main = 2)
abline(h = 12, col = "red")
dev.off()

# exclude outlier
clust = cutreeStatic(sampleTree, cutHeight = 12, minSize = 10) 
table(clust)
keepSamples = (clust==1)
datExpr = datExpr0[keepSamples, ]
nGenes = ncol(datExpr)
nSamples = nrow(datExpr)

nGenes
nSamples

##decide on the softpower threshold; COFFEE BREAK: this could take some time##
powers = c(c(1:10), seq(from = 12, to=20, by=2))
sft = pickSoftThreshold(datExpr, powerVector = powers, verbose = 5)
pdf(file="DNAmLiver_powerplot.pdf", width=9, height=5)
par(mfrow=c(1,2))
cex1 = 0.9
plot(sft$fitIndices[,1], -sign(sft$fitIndices[,3])*sft$fitIndices[,2], xlab="Soft Threshold (power)",ylab="Scale Free Topology Model Fit,signed R^2",type="n", main = paste("Scale independence"))
text(sft$fitIndices[,1], -sign(sft$fitIndices[,3])*sft$fitIndices[,2], labels=powers,cex=cex1,col="red")
abline(h=0.90,col="red")
plot(sft$fitIndices[,1], sft$fitIndices[,5], xlab="Soft Threshold (power)",ylab="Mean Connectivity", type="n", main = paste("Mean connectivity"))
text(sft$fitIndices[,1], sft$fitIndices[,5], labels=powers, cex=cex1,col="red")
dev.off()
# softpower = 6 seems good
softPower=6

# compute adjacencies, and turn that to topological matrix
adjacency = adjacency(datExpr, power = softPower)
# this could take a while; drink more coffee
TOM = TOMsimilarity(adjacency)
dissTOM = 1-TOM

# constructing the CpG trees
geneTree = hclust(as.dist(dissTOM), method = "average")
pdf(file="DNAmLiver_geneTreePower6.pdf", width=12, height=9)
plot(geneTree, xlab="", sub="", main = "Gene clustering on TOM-based dissimilarity", labels = FALSE, hang = 0.04)
dev.off()

# set parameters for modules
minModuleSize = 35
dynamicMods = cutreeDynamic(dendro = geneTree, distM = dissTOM,
              deepSplit = 2, pamRespectsDendro = FALSE,
              minClusterSize = minModuleSize);
table(dynamicMods)

dynamicColors = labels2colors(dynamicMods)
table(dynamicColors)

moduleAndColors = table(dynamicColors, dynamicMods)
# merge modules
MEList = moduleEigengenes(datExpr, colors = dynamicColors)
MEs = MEList$eigengenes
MEDiss = 1-cor(MEs)
METree = hclust(as.dist(MEDiss), method = "average")
MEDissThres = 0.3
pdf(file="DNAmLiver_ME_dedrogram_Power6.pdf", width=12, height=9)
plot(METree, main = "Clustering of module eigengenes", xlab = "", sub = "")
abline(h=MEDissThres, col = "red")
dev.off()

merge = mergeCloseModules(datExpr, dynamicColors, cutHeight = MEDissThres, verbose = 3)
mergedColors = merge$colors
mergedMEs = merge$newMEs
pdf(file="DNAmLiver_geenTree_merged_Power6.pdf", width=12, height=9)
plotDendroAndColors(geneTree, cbind(dynamicColors, mergedColors), c("Dynamic Tree Cut", "Merged dynamic"), dendroLabels = FALSE, hang = 0.03, addGuide = TRUE, guideHang = 0.05)
dev.off()

# inspecting & exporting modules
moduleColors = mergedColors
colorOrder = c("grey", standardColors(50))
moduleLabels = match(moduleColors, colorOrder)-1


Merged_Modules = table(moduleColors, moduleLabels)
dim(Merged_Modules)
write.csv(Merged_Modules, file="DNAmLiver_Module_sizes__Power6.csv")

# can save to file
save(MEs, moduleLabels, moduleColors, geneTree, file = "DNAmLiver_27966CpG_network_Power6.RData")


#Exporting the eivenvalues and modules


# exports dendrogram of the merged modules
MEs0 = moduleEigengenes(datExpr, moduleColors)$eigengenes
MEs = orderMEs(MEs0)
pdf(file="DNAmLiver_moduleMergedCorMatrixSorted_power6.pdf", width=7, height=6)
plotEigengeneNetworks(MEs, "", marDendro = c(0,4,1,2), marHeatmap = c(3,4,1,2), cex.lab = 0.8, xLabelsAngle = 90)
dev.off()

# export inter-module (meta-network) correlations
MEcorMatrix=signif(cor(MEs, use="p"),2)
head(MEcorMatrix)
dim(MEcorMatrix)
write.csv(MEcorMatrix, "DNAmLiver_MEcorrelations_power6.csv")

# saving the module eigengenes as csv
MEall=as.data.frame(MEs)
head(MEall)
dim(MEall)
write.csv(MEall, "DNAmLiver_MEs_power6.csv")

# saving CpG-level connectivities to each module as csv
datKME = signedKME(datExpr, MEs0, outputColumn = "MM")
datKME_colorsLabels=data.frame(datKME, Colors, labels)
write.csv(datKME_colorsLabels, "DNAmLiver_gene-levelKME_power6.csv")

# saving CpG-level intramodule connectivities as csv
Alldegrees1=intramodularConnectivity(adjacency, moduleColors)
moduleConv = data.frame(Alldegrees1)
moduleK_labels_colors=data.frame(moduleConv, Colors, labels)
write.csv(moduleK_labels_colors, "DNAmLiver_moduleKs_power6.csv")
```
