## Supplementary material for "Pleiotropic Influence of DNA Methylation QTLs on physiological and aging traits": All supplemental files: SupTablesFigures_meQTL_10Apr2023.docx

**Supplemental Tables**

**Table S1: Enrichment or depletion in chromatin states and genomic regions among CpGs modulated by meQTLs**

|  |  | **Cis-meQTLs** | | | | **Trans-meQTLs** | | | |
| --- | --- | --- | --- | --- | --- | --- | --- | --- | --- |
| **ChromHMM**  **P0 liver^1^** | **Total in**  **array**  **background** | **Counts** | **Fold**  **enrichment/**  **depletion** | **Enrichment**  **p^2^** | **Depletion**  **p^2^** | **Counts** | **Fold**  **enrichment/**  **depletion** | **Enrichment**  **p^2^** | **Depletion**  **p^2^** |
| En-Pd | 138 | 9 | -1.0 | 0.56 | 0.58 | 25 | 2.4 | **3.0E-05** | 1 |
| En-Pp | 197 | 13 | 1.0 | 0.53 | 0.58 | 24 | 1.6 | 0.01 | 0.99 |
| En-Sd | 56 | 3 | -1.2 | 0.72 | 0.50 | 9 | 2.2 | 0.02 | 0.99 |
| En-Sp | 132 | 13 | 1.5 | 0.09 | 0.95 | 23 | 2.3 | **0.0001** | 1 |
| En-W | 93 | 5 | -1.2 | 0.74 | 0.42 | 13 | 1.9 | 0.02 | 0.99 |
| Hc-H | 33 | 2 | -1.1 | 0.65 | 0.63 | 2 | -1.2 | 0.72 | 0.55 |
| Hc-P | 2948 | 192 | -1.0 | 0.55 | 0.48 | 221 | 1.0 | 0.48 | 0.54 |
| NRS | 5774 | 439 | 1.2 | 0.0002 | 1 | 516 | 1.2 | 1.6E-06 | 1 |
| NS | 6840 | 612 | 1.4 | 3.2E-19 | 1 | 416 | -1.2 | 1 | 1.8E-07 |
| Pr-A | 1572 | 14 | -7.4 | 1 | 2.3E-30 | 69 | -1.7 | 1 | 2.1E-07 |
| Pr-B | 1695 | 56 | -2.0 | 1 | 8.4E-10 | 107 | -1.2 | 0.97 | 0.03 |
| Pr-F | 108 | 10 | 1.4 | 0.17 | 0.90 | 17 | 2.1 | 0.003 | 1 |
| Pr-W | 273 | 9 | -2.0 | 0.99 | 0.01 | 33 | 1.6 | 0.004 | 1 |
| Tr-I | 137 | 12 | 1.3 | 0.19 | 0.89 | 16 | 1.6 | 0.05 | 0.97 |
| Tr-P | 3903 | 300 | 1.2 | 0.001 | 1 | 295 | 1.0 | 0.42 | 0.61 |
| Tr-S | 4067 | 144 | -1.9 | 1 | 1.2E-19 | 302 | -1.0 | 0.55 | 0.47 |
| **Genomic**  **annotation** | **Total in**  **array**  **background** | **Counts** | **Fold**  **enrichment/**  **depletion** | **Enrichment**  **p** | **Depletion**  **p** | **Counts** | **Fold**  **enrichment/**  **depletion** | **Enrichment**  **p** | **Depletion**  **p** |
| intergenic  upstream | 2302 | 247 | 1.6 | 2.7E-15 | 1 | 185 | 1.1 | 0.15 | 0.87 |
| Promoter | 1487 | 55 | -1.8 | 1 | 5.9E-07 | 70 | -1.6 | 1 | 6.7E-06 |
| fiveUTR | 2333 | 72 | -2.1 | 1 | 6.4E-15 | 154 | -1.1 | 0.96 | 0.05 |
| Exon | 9453 | 510 | -1.2 | 1 | 7.8E-09 | 701 | -1.0 | 0.60 | 0.42 |
| Intron | 7348 | 613 | 1.3 | 1.1E-12 | 1 | 590 | 1.1 | 0.02 | 0.98 |
| threeUTR | 2446 | 104 | -1.5 | 1 | 2.3E-07 | 193 | 1.1 | 0.21 | 0.81 |
| intergenic  downstream | 2597 | 232 | 1.4 | 5.1E-07 | 1 | 195 | 1.0 | 0.48 | 0.55 |

^1^15-states chromatin model for P0 mouse liver

^2^Hypergeometric test used for enrichment/depletion. For enrichment, R code used was: phyper(q-1, m, n, k, lower.tail=FALSE). For depletion, R code used was: phyper(q, m, n, k)

**Table S2. Enrichment or depletion in differentially methylated CpGs among CpGs modulated by cis- and trans-meQTLs**

|  |  | **Cis-meQTLs** | | | | **Trans-meQTLs** | | | |
| --- | --- | --- | --- | --- | --- | --- | --- | --- | --- |
| **Main factor** | **Total DMC^1^** | **Counts** | **Fold**  **enrichment/**  **depletion** | **Enrichment**  **p^2^** | **Depletion**  **p^2^** | **Counts** | **Fold**  **enrichment/**  **depletion** | **Enrichment**  **p^2^** | **Depletion**  **p^2^** |
| Age-gain | 5030 | 246 | -1.3 | 1 | 3.2E-08 | 605 | 1.6 | 8.8E-38 | 1 |
| Age-loss | 1523 | 58 | -1.7 | 1.00 | 1.2E-06 | 99 | -1.1 | 0.94 | 0.08 |
| Weight | 733 | 34 | -1.4 | 0.99 | 0.02 | 94 | 1.7 | 1.8E-07 | 1 |
| Diet | 321 | 10 | -2.1 | 1.00 | 0.005 | 55 | 2.3 | 6.0E-09 | 1 |
| Lifespan | 236 | 27 | 1.7 | **0.003** | 1 | 43 | 2.4 | 4.3E-08 | 1 |

^1^Number of differentially methylated CpG at Bonferroni p ≤ 0.05 (as reported in PMID: 35389339)

^2^Hypergeometric test used for enrichment/depletion. For enrichment, R code used was: phyper(q-1, m, n, k, lower.tail=FALSE). For depletion, R code used was: phyper(q, m, n, k)

**Table S3. Markers associated with >20 meQTLs**

|  | **maxLOD marker** | **maxLOD Chr** | **maxLOD Mb** | **Total meQTL** | **trans-meQTL**  **(n)** | **cis-**  **meQTL**  **(n)** | **Hotspot**  **type** | **Modules** | **Broad location** |
| --- | --- | --- | --- | --- | --- | --- | --- | --- | --- |
| 1 | *rs33188980* | 2 | 107.07 | 21 | 2 | 19 | cis |  | *Chr2:102–112* |
| 2 | *rs27248595* | 2 | 144.09 | 30 | 30 | 0 | trans | *Green (2092 CpGs)* | *Chr2:141–151* |
| 3 | *rs27248575* | 2 | 144.09 | 27 | 26 | 1 |  |  |  |
| 4 | *rs29541164* | 2 | 149.64 | 35 | 33 | 2 |  |  |  |
| 5 | *rs6206791* | 2 | 149.67 | 27 | 21 | 6 |  |  |  |
| 6 | *rs27475809* | 4 | 119.38 | 23 | 0 | 23 | cis |  | *Chr4:114-124* |
| 7 | *rs47259742* | 5 | 114.60 | 45 | 39 | 6 | trans, *meQTL.Chr5a* | Blue (5067 CpGs) | *Chr5:110-120* |
| 8 | *rs51209772* | 5 | 114.87 | 59 | 59 | 0 |  |  |  |
| 9 | *rs29733222* | 5 | 115.43 | 233 | 230 | 3 |  |  |  |
| 10 | *rs36365158* | 5 | 116.41 | 81 | 78 | 3 |  |  |  |
| 11 | *rs260237020* | 5 | 116.56 | 49 | 48 | 1 |  |  |  |
| 12 | *rs50259190* | 7 | 137.86 | 35 | 34 | 1 | trans |  | *Chr7:132–142* |
| 13 | *rs3658866* | 14 | 22.96 | 30 | 11 | 19 | cis |  | *Chr14:17-27* |
| 14 | *rs48387725* | 14 | 46.44 | 30 | 29 | 1 | trans |  | *Chr14: 41–51* |
| 15 | *rs46161522* | 14 | 46.98 | 34 | 32 | 2 |  | *Black (but removed when cofactors included)*  *Greenyellow (998 CpGs)* |  |
| 16 | *rs31048374* | 14 | 49.09 | 29 | 27 | 2 |  |  |  |
| 17 | *rs30567369* | 19 | 47.51 | 30 | 0 | 30 | cis | *Royalblue (62 CpGs)* | *Chr19:42–52* |
| 18 | *rs31157694 (royalblue)* | 19 | 47.94 | 29 | 1 | 28 |  |  |  |

^1^This is the nearest gene for intergenic CpGs

^2^This CpG located near *Foxa2* maps as a cis-meQTL and does not map to Chr5; however, other CpGs near *Foxa2* map as *trans-meQTLs* to meQTL.Chr5a.

**Table S4. Enrichment or depletion is chromatin states and genomic regions among CpGs with trans-meQTL in *meQTL.5a***

| **ChromHMM**  **P0 liver** | **Array**  **background** | **meQTL.5a (n)** | **Fold enrichment/depletion** | **Enrichment p** | **Depletion p** |
| --- | --- | --- | --- | --- | --- |
| En-Pd | 138 | 15 | 6.1 | **2.9E-08** | 1.00 |
| En-Pp | 197 | 11 | 3.1 | **0.0009** | 1.00 |
| En-Sd | 56 | 8 | 8.0 | **6.6E-06** | 1.00 |
| En-Sp | 132 | 13 | 5.5 | **7.6E-07** | 1.00 |
| En-W | 93 | 8 | 4.8 | **0.0003** | 1.00 |
| Hc-P | 2948 | 26 | -2.0 | 1.00 | 1.4E-05 |
| NRS | 5774 | 167 | 1.6 | 1.4E-11 | 1 |
| NS | 6840 | 78 | -1.6 | 1.00 | 7.5E-07 |
| Pr-A | 1572 | 12 | -2.3 | 1.00 | 0.0004 |
| Pr-B | 1695 | 28 | -1.1 | 0.70 | 0.38 |
| Pr-F | 108 | 10 | 5.2 | **2.5E-05** | 1.00 |
| Pr-W | 273 | 8 | 1.6 | 0.12 | 0.94 |
| Tr-I | 137 | 10 | 4.1 | **0.0002** | 1.00 |
| Tr-P | 3903 | 53 | -1.3 | 0.99 | 0.01 |
| Tr-S | 4067 | 53 | -1.4 | 1.00 | 0.005 |
| Hc-H | 33 | 0 | 6.1 | 1 | 0.55 |
| **Genomic**  **annotation** | **Array**  **background** | **meQTL.5a (n)** | **Fold enrichment/depletion** | **Enrichment p** | **Depletion p** |
| Intergenic_upstream | 2302 | 31 | -1.3 | 0.96 | 0.05 |
| Promoter | 1487 | 10 | -2.7 | 1.00 | 0.0001 |
| fiveUTR | 2333 | 42 | 1.0 | 0.51 | 0.56 |
| Exon | 9453 | 152 | -1.1 | 0.95 | 0.06 |
| Intron | 7348 | 199 | 1.5 | **2.03E-11** | 1 |
| threeUTR | 2446 | 37 | -1.2 | 0.88 | 0.16 |
| Intergenic_downstream | 2597 | 29 | -1.6 | 1.00 | 0.003 |

^1^15-states chromatin model for P0 mouse liver

^2^Hypergeometric test used for enrichment/depletion. For enrichment, R code used was: phyper(q-1, m, n, k, lower.tail=FALSE). For depletion, R code used was: phyper(q, m, n, k)

**Table S5. Number of nodes connected to candidate genes for trans-modulated CpGs, transcripts, and proteins.**

| **Candidate genes** | **node_degree_CpGs** | **node_degree_mRNA** | **node_degree_protein** |
| --- | --- | --- | --- |
| *Hnf1a* | 13 | 1 | 1 |
| *Pxn* | 12 | 4 | 5 |
| *Srsf9* | 10 | 3 | 0 |
| *Rplp0* | 4 | 12 | 5 |
| *Acads* | 3 | 1 | 2 |
| *Ccdc64* | 2 | 1 | 0 |
| *Coq5* | 2 | 0 | 1 |
| *Git2* | 2 | 3 | 2 |
| *Rab35* | 2 | 3 | 1 |
| *Sirt4* | 2 | 1 | 0 |
| *Cit* | 1 | 1 | 1 |
| *Prkab1* | 1 | 3 | 1 |
| *Triap1* | 1 | 1 | 0 |
| *1500011B03Rik* | 0 | 0 | 0 |
| *Gatc* | 0 | 0 | 0 |
| *Hspb8* | 0 | 2 | 3 |
| *Oasl2* | 0 | 13 | 8 |
| *Tchp* | 0 | 1 | 0 |

**Table S6. Enrichment or depletion in differentially methylated CpGs among among CpGs with trans-meQTL in *meQTL.5a***

| **Main factor** | **Array**  **background** | **meQTL.5a**  **(n)** | **Fold**  **enrichment/depletion** | **Enrichment**  **p** | **Depletion**  **p** |
| --- | --- | --- | --- | --- | --- |
| Age-gain | 5030 | 301 | 3.2 | 5.4E-92 | 1 |
| Age-loss | 1523 | 43 | 1.5 | 0.006 | 1 |
| Weight-DMC | 733 | 33 | 2.4 | 4.4E-06 | 1 |
| Diet-DMC | 321 | 15 | 2.5 | 0.001 | 1 |
| LS-DMC | 236 | 0 | 0 | 1 | 0.01 |

^2^Hypergeometric test used for enrichment/depletion. For enrichment, R code used was: phyper(q-1, m, n, k, lower.tail=FALSE). For depletion, R code used was: phyper(q, m, n, k)

**Supplemental Figures**

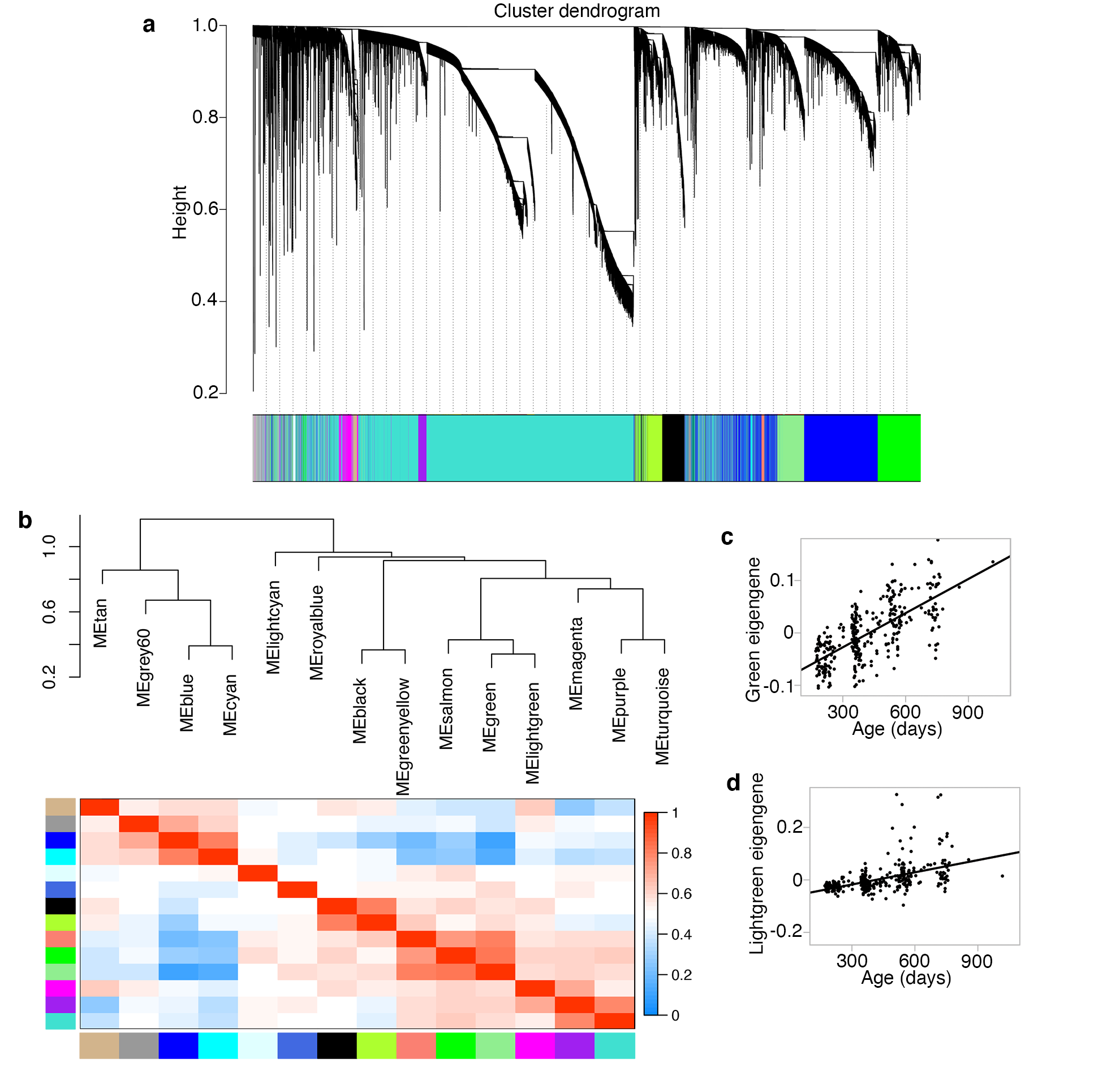

**Fig S1. Weighted gene co-methylation network analysis (WGCNA)**

**(a)** The dendrogram shows the clustering of the CpGs into co-methylation modules. **(b)** Hierarchical clustering of the CpG modules based on correlation between the module eigengenes (ME). Correlation between age and the ME for the **(c)** Green, and **(d)** Lightgreen modules.

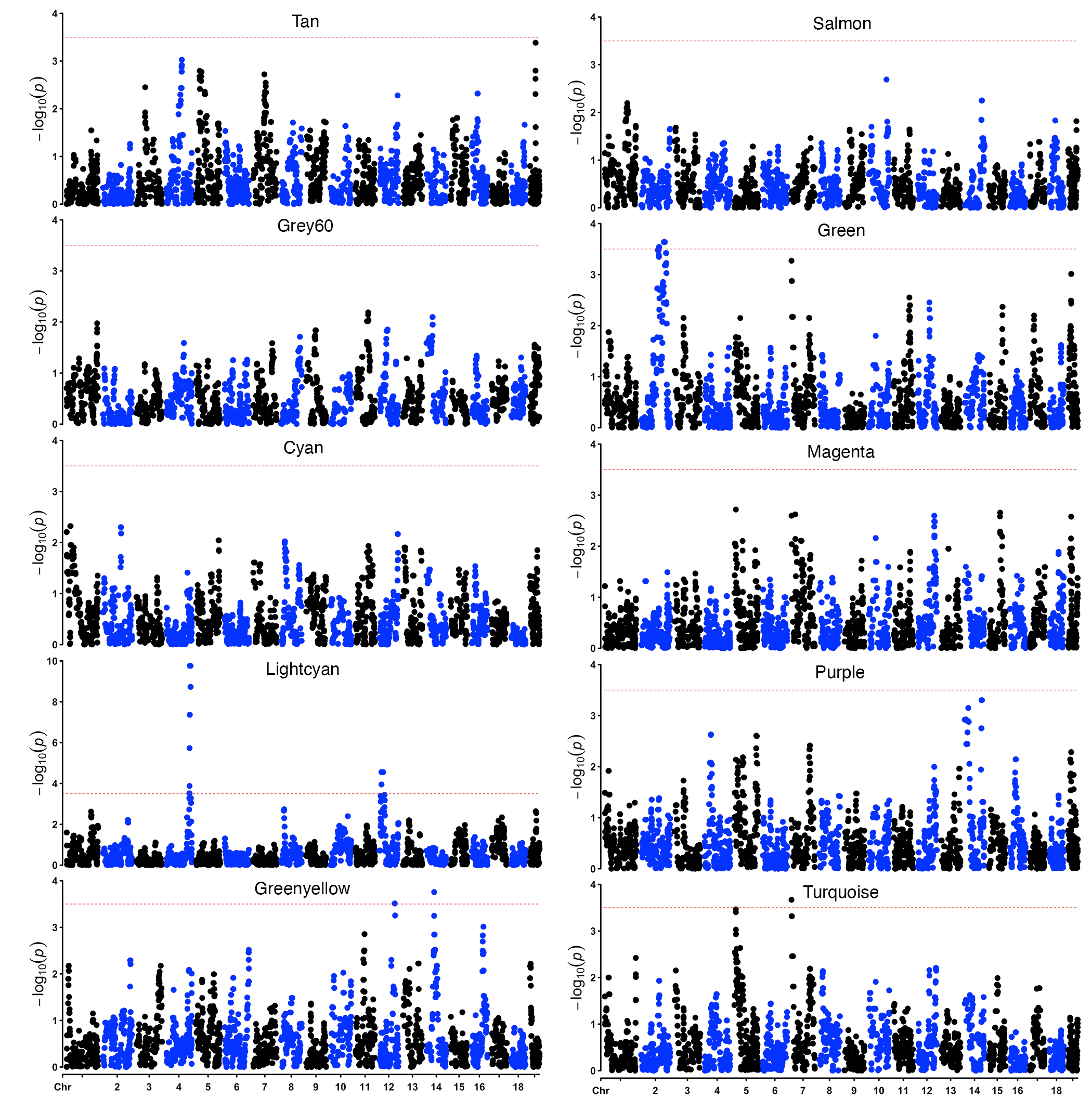

**Fig S2. Genetics of co-methylation CpG networks**

QTL maps for the module eigengenes (MEs). Mapping was done using a linear mix model with adjustment for age, diet, and body weight. The horizontal dashed red line marks a relatively lenient threshold of –log_10_p = 3.5.

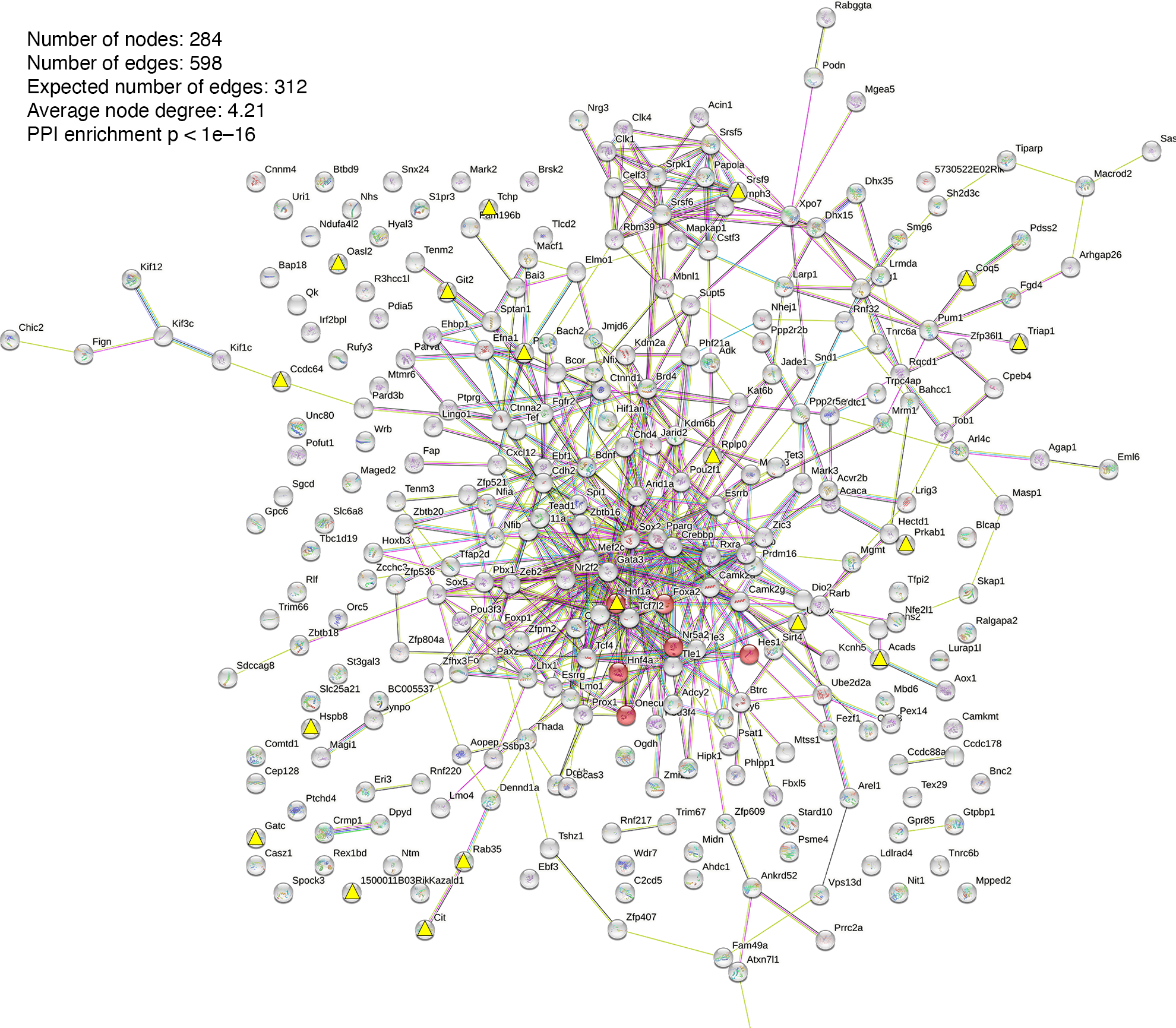

**Fig S2. Protein-protein interaction networks based on trans-meQTLs and candidate genes.** Yellow triangles indicate candidate genes located in meQTL.5a. Red nodes are members of the maturity onset diabetes of the young MODY) pathway and includes the candidate gene HNF1A.

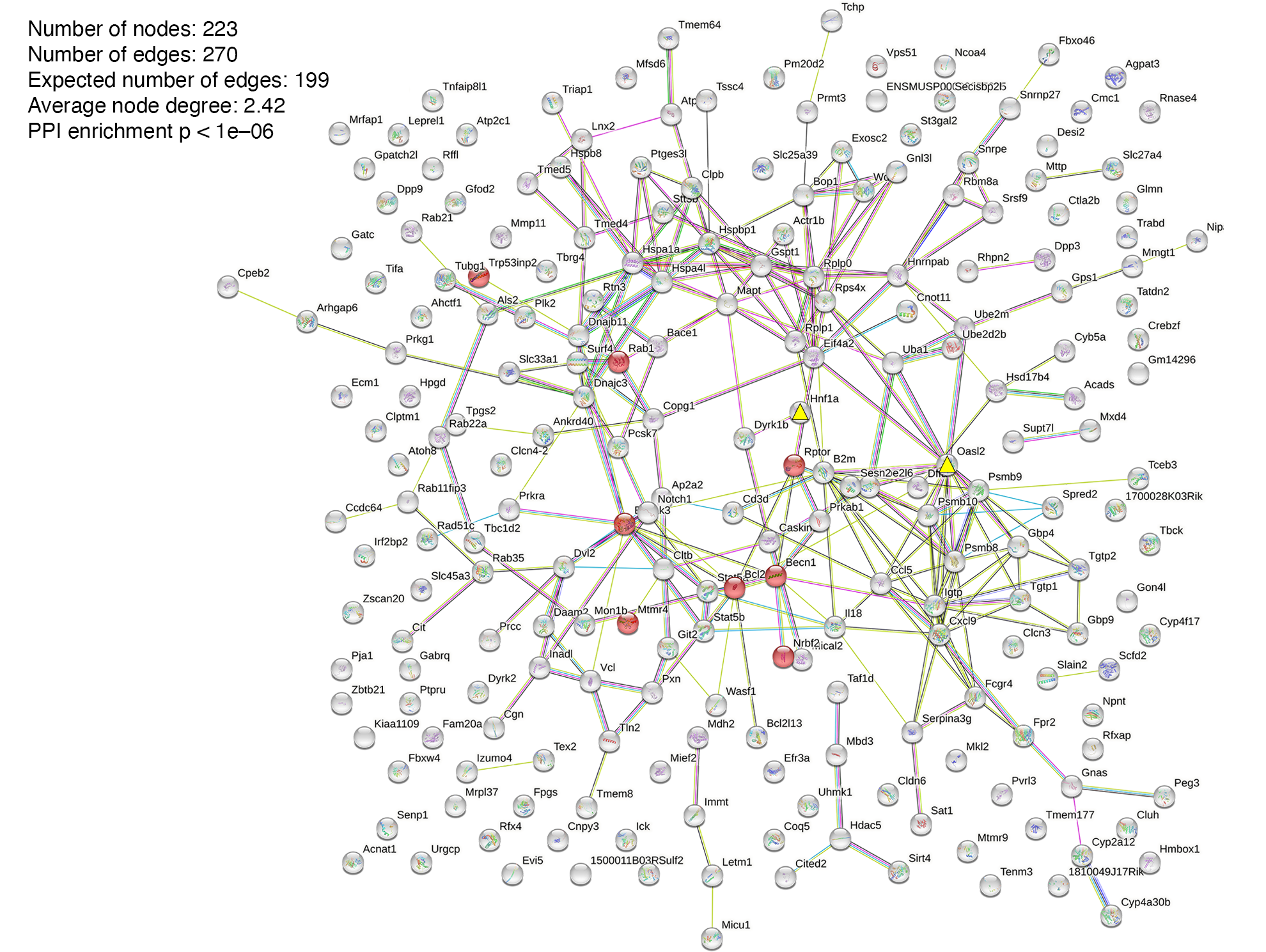

**Fig S4. Protein-protein interaction networks based on trans-eQTLs and candidate genes.**

The yellow triangles denote the location of positional candidates: HNF1A and OASL2. HNF1A was the most highly connected candidate in the meQTL based-network but has low degree of connections with the trans-eQTLs. Among the candidates, OASL2 has the highest number of connections with the trans-eQTLs.

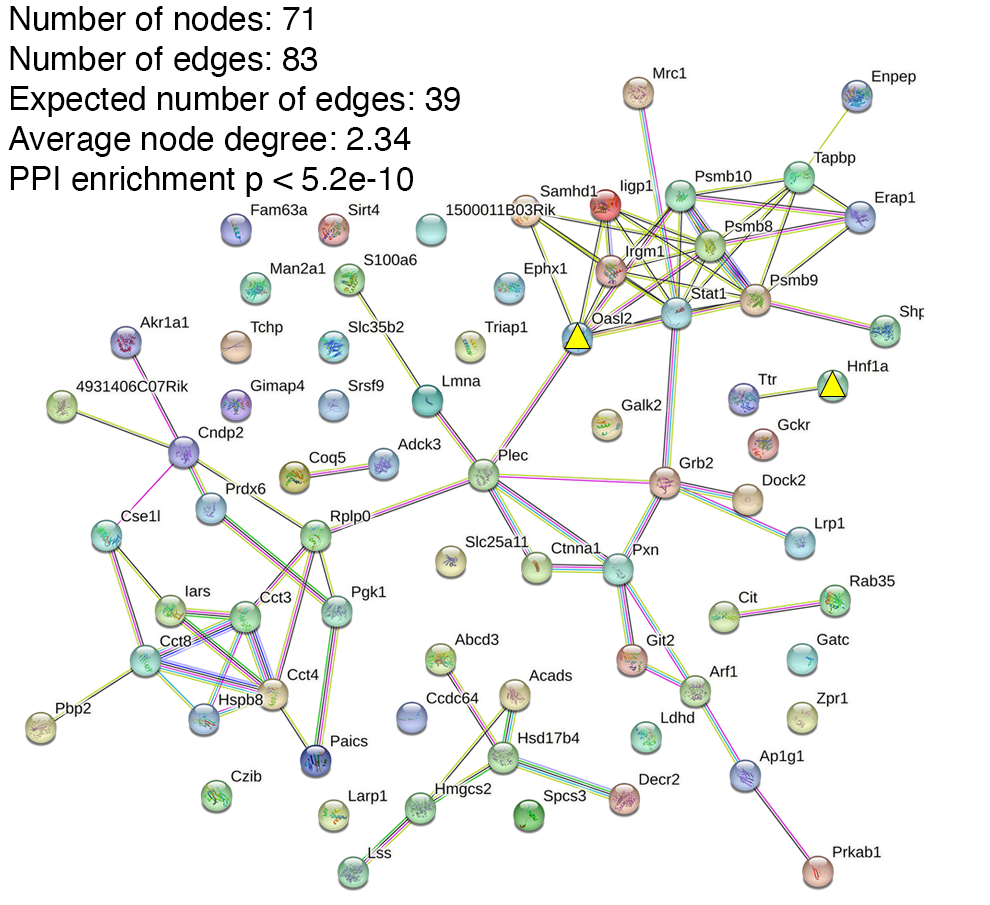

**Fig S5. Protein-protein interaction networks based on trans-pQTLs and candidate genes.**

The yellow triangles denote the location of positional candidates: HNF1A and OASL2. HNF1A was the most highly connected candidate in the meQTL based-network but has low degree of connections with the trans-pQTLs. Among the candidates, OASL2 has the highest number of connections with the trans-pQTLs.

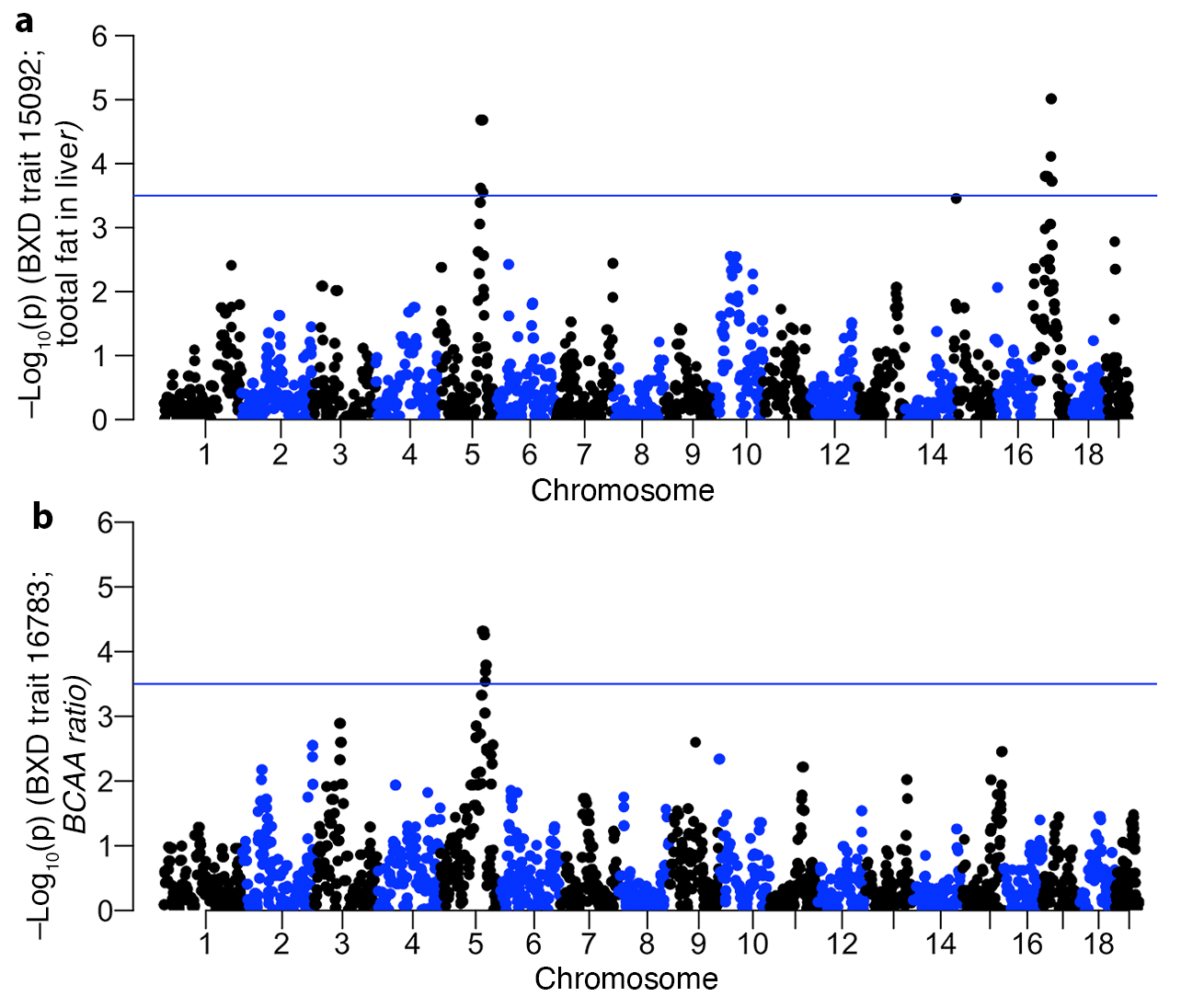

**Fig S6. BXD Metabolic traits linked to meQTL.5a**

QTL maps for **(a)** total fat in liver, and **(b)** ratio of branched chain amino acids to total amino acids. Mapping was done using a linear mix model. The horizontal line marks a relatively lenient threshold of –log_10_p = 3.5.
